# Expression of the human immunomodulatory protein, human B7-1 (CD80), accelerates neuroinflammation, synaptic loss, microvascular instability and lethality in a murine model of Alzheimer’s Disease

**DOI:** 10.64898/2025.12.30.697078

**Authors:** Victor Danelon, Sarah C. Garrett-Thomson, Nicholas C. Morano, Ruben Constanza, Pouneh Kermani, Roshelle S. Smith, Steven C Almo, Francis S. Lee, Barbara L. Hempstead

## Abstract

Immune-mediated inflammatory processes play a pivotal role in the pathogenesis of Alzheimer’s disease (AD). However, immune loci exhibit significant sequence diversity, with human proteins sharing only ∼45–70% identity with their murine orthologs. This divergence contributes to the inability of many established mouse models to accurately capture key neuroinflammatory mechanisms relevant to human AD. We recently identified that the human, but not murine, immunomodulatory protein B7-1 (CD80) activates the p75 neurotrophin receptor (p75), a function arising from evolutionary divergence in human B7-1. This discovery provides an opportunity to directly interrogate the role of this interaction in disease progression using a well-characterized murine model of mutant Aβ overexpression (CRND8). We generated a mouse line in which murine B7-1 was replaced with a chimeric human:murine B7-1 that retains normal interactions with CTLA-4 and CD28, while gaining the ability to bind p75, and evaluated its effects in CRND8 mice. Expression of human:murine B7-1 in vivo resulted in increased lethality, accelerated neuroinflammation of resident glia, more rapid synaptic and dendritic loss, and enhanced microvascular compromise in the subiculum compared to CRND8 mice expressing murine B7-1. Together, these findings identify the human B7-1:p75 interaction as a previously unrecognized contributor to AD pathogenesis and a potential therapeutic target in a brain region critical for learning and memory that is affected in early stages of disease.

## Introduction

Alzheimer’s disease (AD) is the leading cause of cognitive dysfunction, with progressive memory deficits causing abnormalities in spatial processing, language and judgement that severely compromise daily living (reviewed in (Knopman et al 2021)). AD is characterized by neuropathological lesions of amyloid plaques, which consist of misfolded and aggregated amyloid-β (Aβ) in the extracellular space, as well as intra-neuronal neurofibrillary tangles composed of hyperphosphorylated tau. In addition, Aβ accumulation in the vasculature, leading to cerebral amyloid angiopathy, contributes to the neurovascular pathology of aging, which is the leading contributor to cognitive impairment in the extremely old (age greater than 90 years) (reviewed in (Santisteban & Iadecola 2025)). While the preponderance of research has focused on the pathological mechanisms related to abnormal amyloid deposition and tau-induced neurofibrillary tangles, increasing evidence points to a pivotal role for immune processes in the pathogenesis of AD and neurovascular disease (reviewed in (Heneka et al 2025)). Innate immune signaling associated with Aβ plaques involves activated microglia and reactive astrocytes early in the course of disease, which facilitate the release of immune modulators (i.e., cytokines, chemokines) (Heneka et al 2025, Moonen et al 2023, Rogers et al 1988). In early AD, anti-inflammatory subsets of microglia surround Aβ plaques and support clearance by phagocytosis (Boche & Nicoll 2020, Zotova et al 2013). However, as disease progresses, pro-inflammatory subsets of microglia continuously respond to signals released by dying neurons and Aβ plaques, leading to hyperactivated microglia and chronic inflammatory responses (Andrews et al 2023, Grubman et al 2021). Similarly, in early AD, phenotypically normal astrocytes can play a protective role, supporting neuronal function and Aβ clearance, and as the disease progresses, aberrant overstimulation via pro-inflammatory signals results in the transformation of astrocytes into various pathological phenotypes (A1 reactive, Death, Senescence, and Functional Impairment), which all exacerbate the AD phenotype via different mechanisms (Kim et al 2024, Zhou et al 2020). Ultimately, chronic inflammatory signaling in the brain leads to pathological recruitment of bone marrow derived monocytes and macrophage to the brain parenchyma (Thome et al 2018). The reliance on murine models of AD poses a frequently underrecognized challenge due to the significant evolutionary divergence in murine and primate immune systems (Yang et al 2022). Indeed, immune loci are among the most genetically diverse, with human immune lineage proteins sharing only 45-70% identify with their murine orthologs (West & Deng 2019). In diseases that develop as a consequence of maladaptive immune responses, the “illusion of similarity” between the murine and primate immune interactions may mask species-specific differences and contribute to the inability of animal models to fully recapitulate the mechanistic features of human diseases.

The recently described neuroimmune interaction between B7-1 (CD80) and the p75 neurotrophin receptor is restricted to humans and other primates and does not occur in mice and lower mammals (Morano et al 2022). B7-1, a type one transmembrane protein within the immunoglobulin superfamily (IgSF), is composed of extracellular membrane distal IgV and membrane proximal IgC domains. B7-1 is expressed by dendritic cells, macrophages, microglia, reactive astrocytes and B cells, and regulates T cell function through interactions with CD28 and CTLA-4 and PD-L1 (Sugiura et al 2019). B7-1 co-stimulates T cells through interactions with CD28 and co-inhibits T cells through interaction with CTLA-4, actions that have been extensively characterized (Schildberg et al 2016). B7-1 expression is tightly regulated, and its expression is induced by inflammatory pathways in the setting of infection, injury or aging (Ceeraz, Nowak & Noelle 2013). Upregulation of B7-1 on antigen presenting cells occurs in response to numerous inflammatory stimuli, specifically following IFNγ exposure, activation by Toll-like receptors and by infiltrating T helper cells, and is well documented in rodent models of experimental autoimmune encephalitis, stroke (Felger et al 2010) and traumatic brain injury (Turtzo et al 2014). In humans, B7-1 protein is upregulated in multiple sclerosis patients (Genc, Dona & Reder 1997, Windhagen et al 1995) and RNAseq data supports induction in monocytes, astrocytes and microglial subpopulations in brains of patients with AD (Mathys et al 2024). The p75 receptor, a member of the TNFR superfamily (TNFR16) is expressed primarily on limited subpopulations of adult neurons and has been best characterized as a receptor for mature neurotrophins and proneurotrophins (Malik et al 2021). p75 possesses a cytosolic death domain that induces apoptotic signaling, and can destabilize the neuronal cytoskeleton, leading to acute spine elimination (Deinhardt et al 2011, Giza et al 2018, Yamashita, Tucker & Barde 1999). In the adult brain, p75 expression is normally confined to specific regions, but in response to inflammation, injury (Irmady et al 2014, Kokaia et al 1998, Meeker et al 2012, Sebastiani et al 2015), or neurodegenerative diseases such as AD (Coulson 2006), it becomes upregulated in many classes of neurons, impairing their function.

Primate B7-1, but not B7-1 from lower mammals such as mice (with 46% sequence identity between mouse and human) binds to p75, suggesting that recent evolutionary substitutions in the ectodomain of primate B7-1 confer binding (Morano et al 2022). Human B7-1 (hB7-1), but not murine B7-1 (mB7-1), binds to both human p75 and murine p75, consistent with the high evolutionary conservation of p75 (91.8% sequence identity). Initial studies demonstrated that application of hB7-1 to cultured hippocampal neurons expressing endogenous p75 on their postsynaptic spines resulted in altered dendritic morphology, with loss the post synaptic marker PSD95 in a p75-dependent manner. Injection of hB7-1 into the murine subiculum, a hippocampal region that expresses endogenous p75 and is affected in AD, resulted in acute p75-dependent pruning of dendritic spines. These findings indicate that the interaction between hB7-1, a typical ligand on immune cells, and p75 on neurons may play a previously unrecognized role in neuronal dysfunction (Morano et al,2022). This interaction represents an example of functional cross-talk between the immunoglobulin and TNF receptor superfamilies and expands the functional roles of immune cells in the central nervous system. To critically evaluate the impact of endogenously expressed hB7-1 in the neuroinflammatory setting of mutant Aβ overexpression, a widely employed model of AD, we generated a murine knock-in model expressing chimeric h:mB7-1 (i.e., human IgV domain and murine IgC and transmembrane domains) and confirmed that the h:mB7-1 protein binds p75. These studies revealed previously unrecognized effects of h:mB7-1 on amyloid plaque accumulation, the timing of activation of astrocytes and microglia, neuronal degeneration involving spine loss and dendritic destabilization, and microvascular compromise. The effects of hB7-1 on multiple cell types may represent early and targetable contributions to the development of AD.

## Results

### Development of a knock-in mouse model expressing a chimeric human–mouse B7-1 to probe hB7-1–p75–mediated signaling

To study the effects of hB7-1 expression in mice, we generated a knock-in mouse model expressing a chimeric human:mouse B7-1 (h:mB7-1). We evaluated the alignment of human and murine B7-1 in the context of the p75 interacting sites on B7-1 determined by alanine scanning mutagenesis (Morano et al 2022). The N-terminal IgV domain of B7-1 contains the amino acids that confer p75 binding and diverge between the human and murine sequence. The IgV domains are encoded in a single exon, permitting a potential exon swap, substituting the human IgV for the murine IgV domain (Fig. 1A,B). To ensure that this strategy yielded a stable h:mB7-1 chimeric protein that interacted with p75, we expressed the proposed chimera, in which the entire murine IgV segment (106 aa) was replaced by the human sequence (105 aa) sequence, as a C-term GFP fusion protein in mammalian cells. Incubating these cells with recombinant murine p75--Fc protein demonstrated that the h:mB7-1 chimera exhibited p75 binding that was similar to human B7-1. Furthermore, titrations with recombinant mouse CTLA-4-hIgG1 and mouse CD28-hIgG1 (Fig. 1A and Supplemental Fig. 1) demonstrated that the chimeric h:mB7-1 protein bound these receptors similarly to both human B7-1 and mouse B7-1. Based on these results, a chimeric knock-in mouse (h:mB7-1 KI) was generated using CRISPR-based technologies on a C57BL/6J background (Fig. 1B,C).

**Figure 1.**
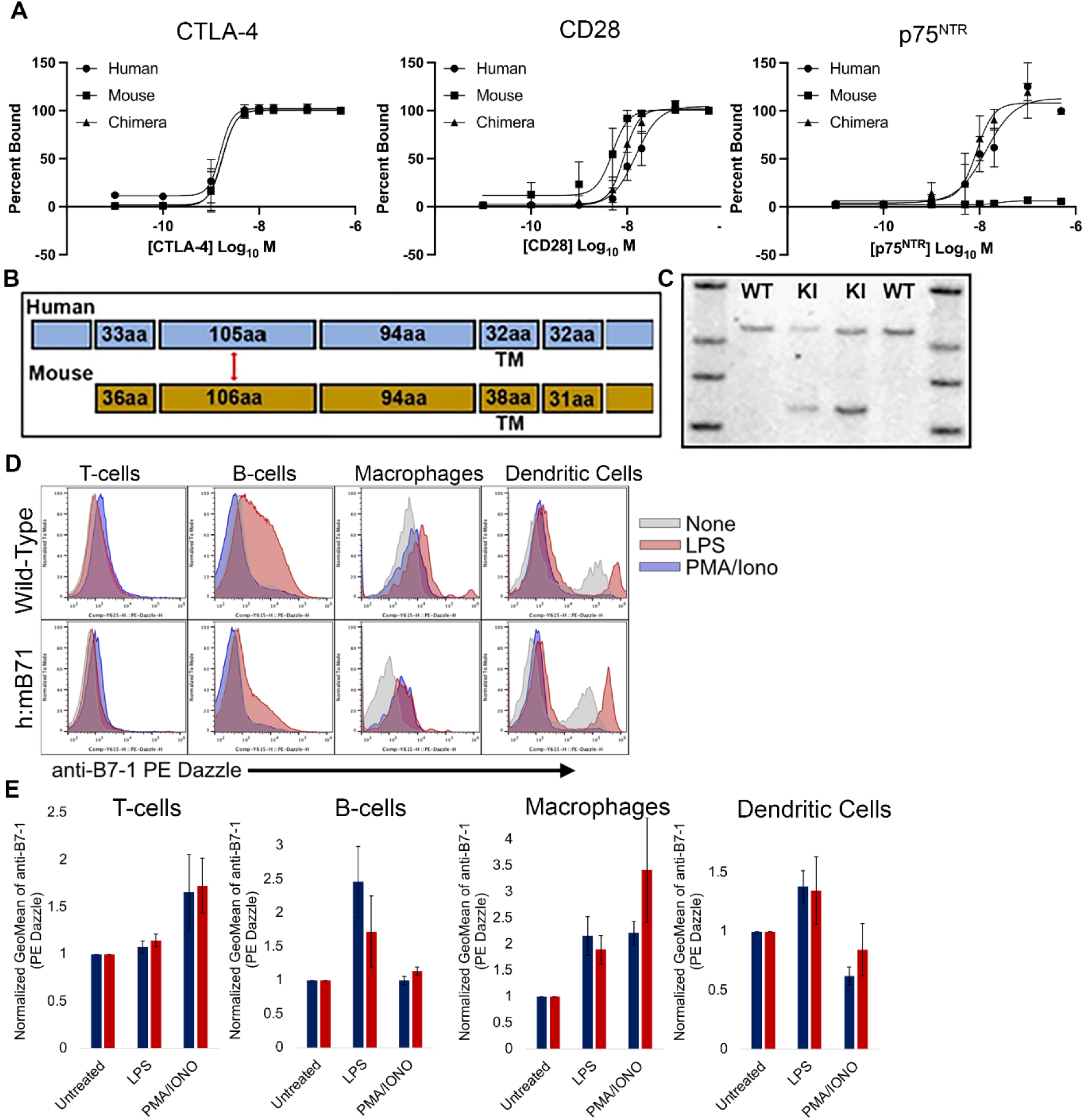
*Development and characterization of a hB7-1 chimera mouse*. A) HEK 293 cells transiently transfected with constructs for either HUMAN, MOUSE, or CHIMERIC exon-swapped B7-1-GFP. Two days post transfection cells were titrated with increasing concentrations of mouse CTLA-4, CD28, and p75^NTR^ hIgG1 fc-fusion proteins and subsequently treated with anti-human A647 antibody. The percent binding was determined by flow cytometry gated as the percentage of GFP positive cells that were also Alexa647 positive (double positive). Note that mp75^NTR^ does not bind human B7-1 expressing cells within the range of concentrations tested. B) IgV domain exon swapping strategy to generate the h:mB7-1 KI mouse. C) Southern blot showing specific human DNA fragment in h:mB7-1 chimeric mice. D and E) Splenocytes were isolated from WT and h:mB71 chimera animals using established methods and stimulated with either LPS or PMA/Ionomycin for 24 hours. After stimulation, cells were collected and stained with cell-surface markers for different immune populations (αmCD45, αmCD3, αmCD19, αmCD11b, αmF4/80, αmCD11c) and human or mouse B7-1. D) Shows flow cytometry histogram plots for B7-1 expression in immune populations as indicated. T-cells were gated as (Singlets, Live, CD45+, CD3+, CD19-), B-cells were gated as (Singlets, Live, CD45+, CD3-, CD19+), macrophages were gated as (Singlets, Live, CD45+, CD3-, CD19-, CD11b+, F4/80+), and dendritic cells were gated as (Singlets, Live, CD45+, CD3-, CD19-, CD11b-, F4/80-, CD11c+). (E) Data shows the average fold-increase in B7-1 expression, compared to untreated populations, upon treatment with LPS or PMA/Ionomycin for wt (N=7) and h:mB7-1 (N= 8) mice.

To evaluate the expression of the h:mB7-1 chimeric protein, we focused on immune cells, recognizing that human and murine B7-1 are upregulated on monocytes and lymphocytes in response to immune activating stimuli. Splenocytes were harvested from wt (wild-type) or h:mB7-1 KI mice and treated with lipopolysaccaharide (LPS) or a combination of phorbol 12-myristate 13-acetate and ionomycin (PMA/Ionomycin), both potent drivers of antigen independent activation. We then analyzed murine B7-1 or h:mB7-1 expression in different immune subpopulations by flow cytometry. Each stimulus induced comparable increases of mB7-1 expression in wt mice and h:mB7-1 expression in chimeric mice in monocyte and lymphocyte populations (Fig. 1 D,E), supporting the use of these engineered mice to study hB7-1:p75 biology.

Next, we compared the expression of h:mB7-1 and mB7-1 proteins in the brains of h:mB7-1-KI and wt mice, respectively (Fig. 2 and Supplementary Fig. 2). Consistent with RNA and protein expression analysis (Serot, 2000; Quintana, 2015; Dando, 2016; Yang, 2022; Mathys, 2024), the levels of h:mB7-1 and mB7-1 protein expression are very low in the brain parenchyma of young adult (3 mo) mice as assessed by immunofluorescence microscopy (Fig. 2A).

**Figure 2.**
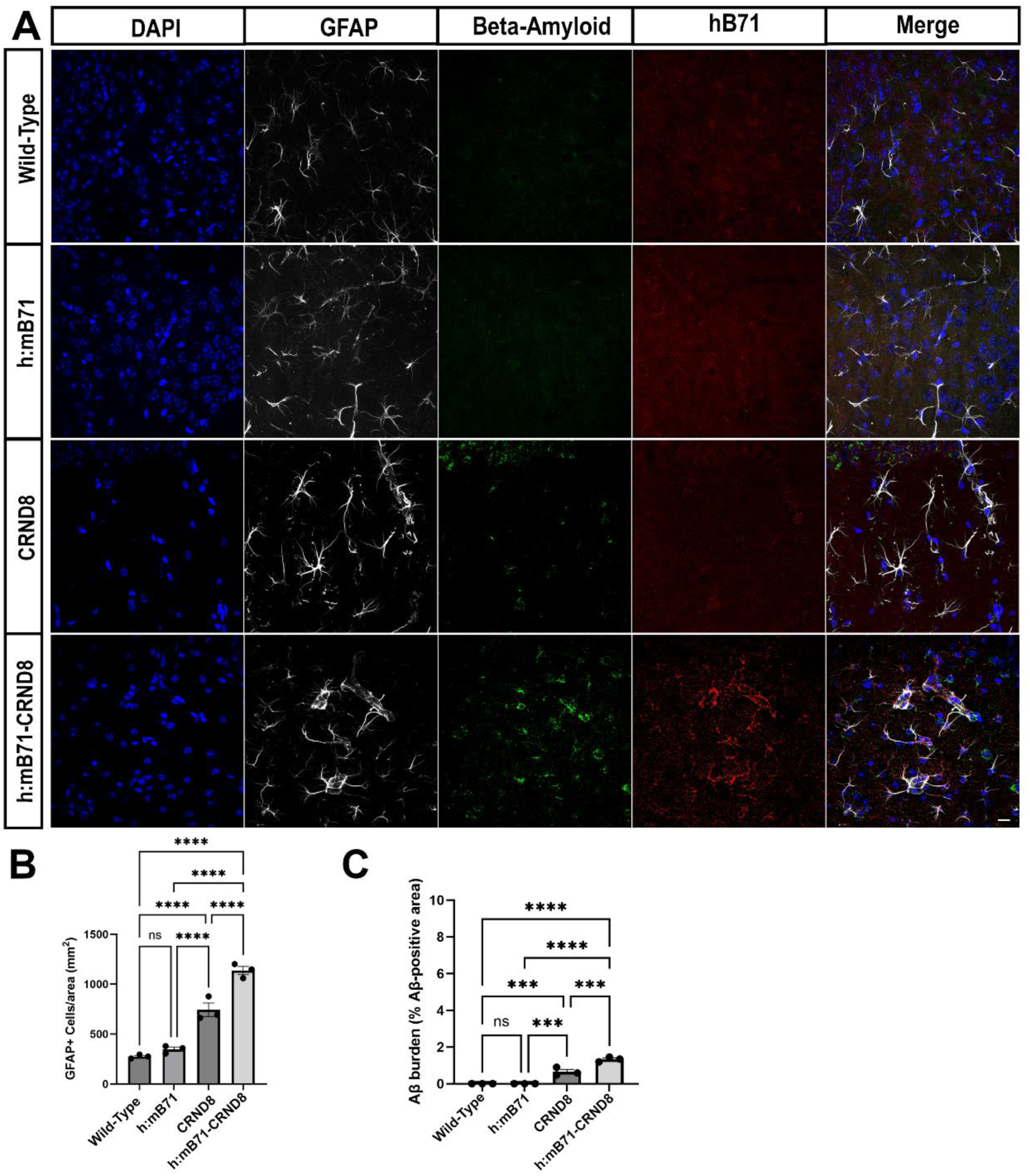
*h:mB71/CRND8 mice have more Aβ burden, GFAP+ cells and hB71 upregulation in the dSubiculum as compared to wt/CRND8 at 3mo*. (A) Representative confocal images of wt/wt, h:mB71/wt, wt/CRND8 and h:mB71/CRND8 mice showing DAPI stained nuclei, GFAP+ cells, βamyloid deposition and hB71 upregulation. hB71 protein is detected only in h:mB71/CRND8 mice. (B) the h:mB71/CRND8 mice have more GFAP+ cells/area and (C) Aβ burden (% positive area) compared to wt/CRND8 mice, wt/wt and h:mB71/wt mice. Wt/CRND8 mice also show significantly more Aβ burden and GFAP+ cells/area as compared to wt/wt and h:mB71/wt. Data shown are mean +/− SEM. In (B)****p>0.0001. In (C) ****p<0.0001, Between CRND8 and hB71-CRND8, ***p=0.0008; between h:mB71, WT to CRND8 ***p=0.0010. (one-way ANOVA followed by post hoc Tukey test). N=3. Scale bars: 10µm.

### Expression of h:mB7-1 accelerates neuroinflammation in an AD model

To investigate the potential roles of hB7-1 in the setting of clinically relevant models of Alzheimer’s Disease, we utilized the CRND8 (hybrid C3H/He-C57BL/6 background) transgenic mouse, which overexpresses mutant human amyloid precursor protein at levels approximately 5 fold higher than endogenous APP (Bellucci et al 2006, Chishti et al 2001). This is an established and very well-characterized transgenic AD model that exhibits rapid accumulation of amyloid plaques, inflammation with astrocyte and microglial activation, synaptic loss and impairments in learning tasks by 3 months of age (Chishti et al 2001, Dudal et al 2004, Jolas et al 2002). Crosses of h:mB7-1/mB7-1 (h:mB7-1/wt) mice with mB7-1/CRND8 (wt/CRND8) mice yielded offspring of 4 genotypes (wt/wt, h:mB7-1/wt, wt/CRND8 and h:mB7-1/CRND8 mice) which were evaluated for expression of B7-1 and p75 proteins. Mice of 2 to 4 months of age were chosen for study, corresponding to conditions prior to onset of Aβ deposition (2 mo), established plaque deposition and microglial and astrocytic activation (3 mo) and severe plaque deposition and astrocyte/microglial clustering around plaques (4 mo). These points reflect the well-established behavior phenotypes of memory impairment at 3 months, and the rapidly progressive nature of disease with lethality of a significant proportion of mice by 4 months of age, an outcome that is modulated by genetic background (Chishti et al 2001, Dudal et al 2004). Prior studies indicated sexual dimorphism in the survival of CRND8 mice (C57BL/l6 background), with approximately 28% of CRND8+ females, and 50% of CRND8+ males dying at three months of age, despite similar levels of Aβ burden as evaluated by Western blot analysis and the density of hippocampal plaques (Cortes-Canteli et al 2019, Granger et al 2016). We compared the natural survival rates, i.e., without additional inflammatory stimuli, of the four genotypes (C57Bl6/CEH-He background) over a 4 month period (Fig. 3). The survival of male and female wt/wt and h:mB7-1/wt mice was greater than 95% at three months of age, indicating that expression of the h:mB7-1 protein in the absence of an inflammatory insult did not impact survival. In contrast, at four months of age, male wt/CRND8 and male h:mB7-1/CRND8 cohorts experienced 29% and 74% lethality, respectively. Sexual dimorphism was also apparent, with 17% lethality of female wt/CRND8 mice, but 30% lethality in female h:mB7-1/CRND8 at four months of age. These data suggest that the expression of h:mB7-1 protein, in the context of mutant Aβ overexpression, results in enhanced lethality. We utilized male mice for all subsequent studies due to the sexual dimorphism in phenotype, and the opportunity to compare our findings directly with prior publications, which exclusively evaluated male CRND8 mice.

**Figure 3.**
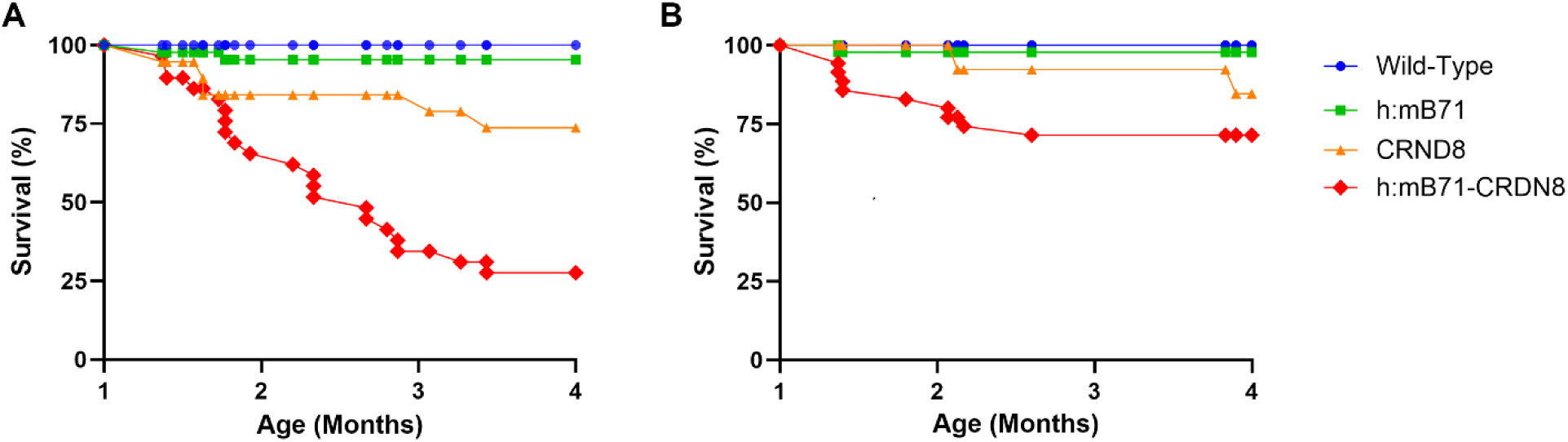
*Survival is impaired in h:mB71/CRND8 mice as compared to wt/CRND8 mice.* Cumulative survival (%) was monitored from 1-4 month of age. (A) In males, the presence of h:mB71 in the context of neuroinflammation induced by AD, results in increased lethality in h:mB71/CRND8 male mice as compared to male mice expressing murine B7-1 (wt/CRND8) and more than wt/wt and h:mB71/wt mice. (B) In females, lethality of h:mB71/CRND8 mice was increased as compared to wt/CRND8 mice and wt/wt and h:mB71/wt mice.

The subiculum is a region of the hippocampal formation that is critically involved in memory formation and retrieval (Ding 2013, Roy et al 2017) and atrophy and degeneration of subicular pyramidal neurons is an indicator of the onset of AD (Carlesimo et al 2015). Importantly, these neurons express p75 during adulthood (Morano et al 2022). Multiple murine models of Aβ overexpression document that the subiculum is affected early, with the density of β-amyloid, GFAP labeling and Iba1 labeling that is 50% greater than other hippocampal subregions (Marongiu et al 2025, Platholi et al 2023). Indeed, prior studies (Morano et al 2022) demonstrated that direct injection of hB7-1-Fc into the subiculum of wild-type mice (3mo) induced rapid (2-14h) pruning of dendritic spines, indicating that these neurons were susceptible to hB7-1 mediated effects, and that this response was dependent on endogenous p75 expression as mice lacking p75 (p75^-/-)^ did not exhibit spine loss (Morano et al 2022).

To determine whether h:mB7-1 and mB7-1 are induced in the subiculum in the setting of Aβ overexpression (CRND8, expressing a mutant human transgene), littermate mice of the 4 genotypes were evaluated at 2, 3 and 4 months of age. At baseline (2mo), expression of murine B7-1, or h:mB7-1 protein was very low in wt mice, as β-amyloid deposition was very low (Supplementary Fig. 4 and data not shown). At 3 months of age, h:mB7-1 protein or murine B7-1 protein remained undetectable in mice lacking the human mutant transgene (Fig. 2). However, in wt/CRND8 mice, murine B7-1 was detectable and some murine B7-1+ cells also expressed GFAP (Supplementary Fig. 2). In h:mB7-1/CRND8 mice, which have one one h:mB7-1 allele and one murine B7-1 allele, the chimeric h:mB7-1 protein and the murine B7-1 protein were induced and detectable in regions where β-amyloid was also expressed, and some of the hB7-1(+) cells co-stained for GFAP (Fig. 2 and Supplementary Fig. 2). By 4 months of age, murine B7-1 and h:mB71 remained undetectable in mice lacking the human mutant transgene (Fig. 4 and Supplementary Fig. 3). However, in CRND8+ mice, murine B7-1 (wt/CRND8 mice), and human and murine B7-1 (h:mB7-1/CRND8 mice) were detectable in regions adjacent to β-amyloid deposition, and some of these cells co-localized with GFAP (Fig. 4 and Supplementary Fig. 3).

**Figure 4.**
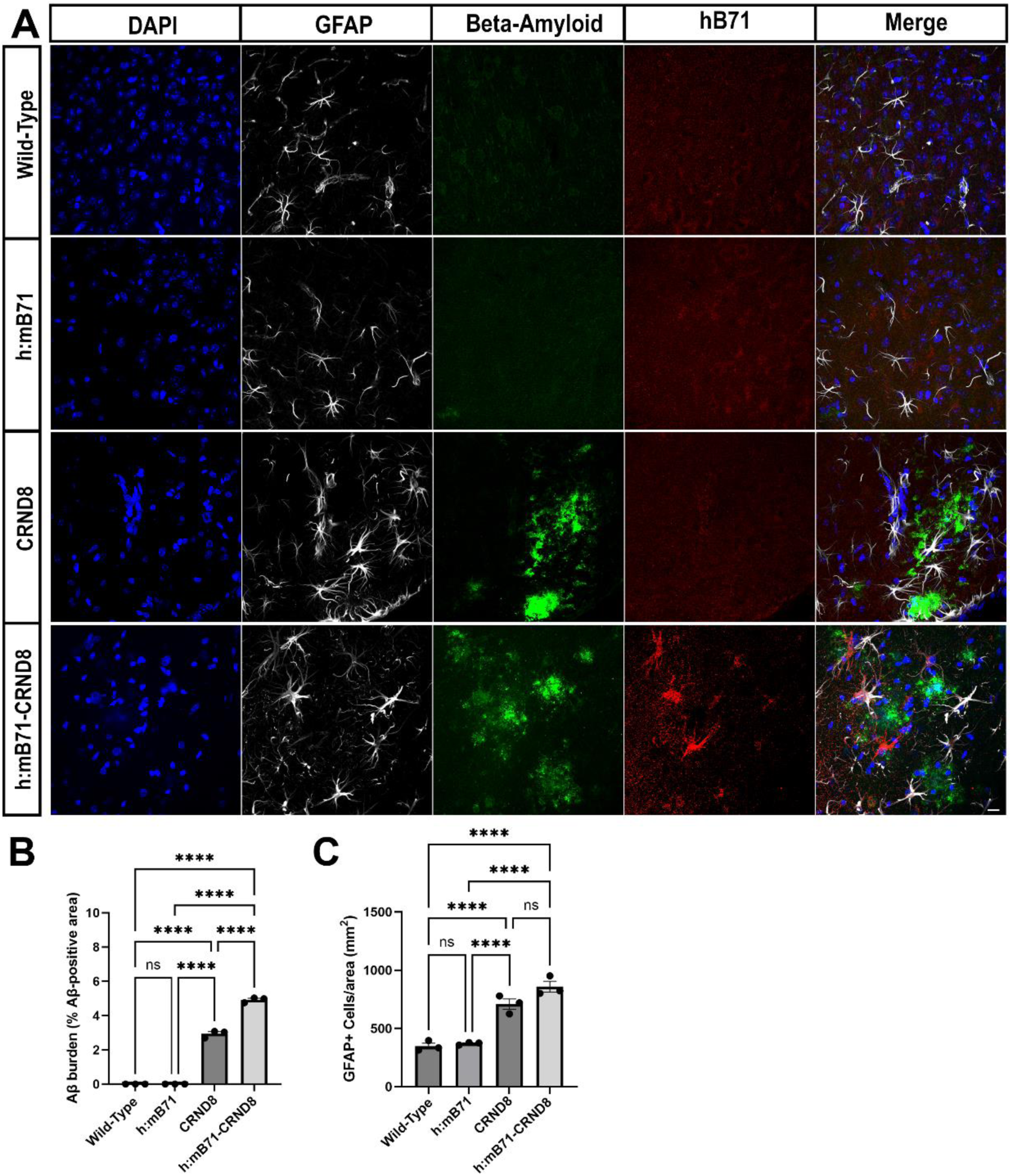
*h:mB71/CRND8 mice hB71 upregulation, as well as more Aβ burden, GFAP+ cells and in the dSubiculum as compared to wt/CRND8 at 4mo.* (A) Representative confocal images of wt/wt, h:mB71/wt, wt/CRND8 and h:mB71/CRND8 mice showing DAPI stained nuclei, GFAP+ cells, βamyloid deposition and hB71 upregulation. hB71 protein is detected only in h:mB71/CRND8 mice. (B) the h:mB71/CRND8 mice have more Aβ burden (% positive area) and (C) GFAP+ cells/area compared to wt/CRND8 mice, wt/wt and h:mB71/wt mice. Wt/CRND8 mice also show significantly more Aβ burden and GFAP+ cells/area compared to wt/wt and h:mB71/wt mice. Data shown are mean +/− SEM. In (B)****p<0.0001. In (C) ****p<0.0001, n.s (p=0.1443) not significant. One-way ANOVA followed by post hoc Tukey test. N=3. Scale bars: 10µm

In the subiculum, expression of p75 in wt and h:mB7-1 expressing mice lacking the CRND8 transgene was limited to fibers, as previously described (Morano et al 2022). At 3 months of age, expression of p75 was induced in fibers of the wt/CRND8 mice (2 fold compared to wt/wt) and was even more prominently expressed in h:mB7-1/CRND8 mice (5 fold compared to wt/wt) (Fig. 5). At 4 months of age this differential elevation of p75 in the h:mB7-1/CRND8 mice (5 fold increase compared to wt animals) and in wt/CRND8 mice (2 fold induction) was maintained (Fig. 6).

**Figure 5.**
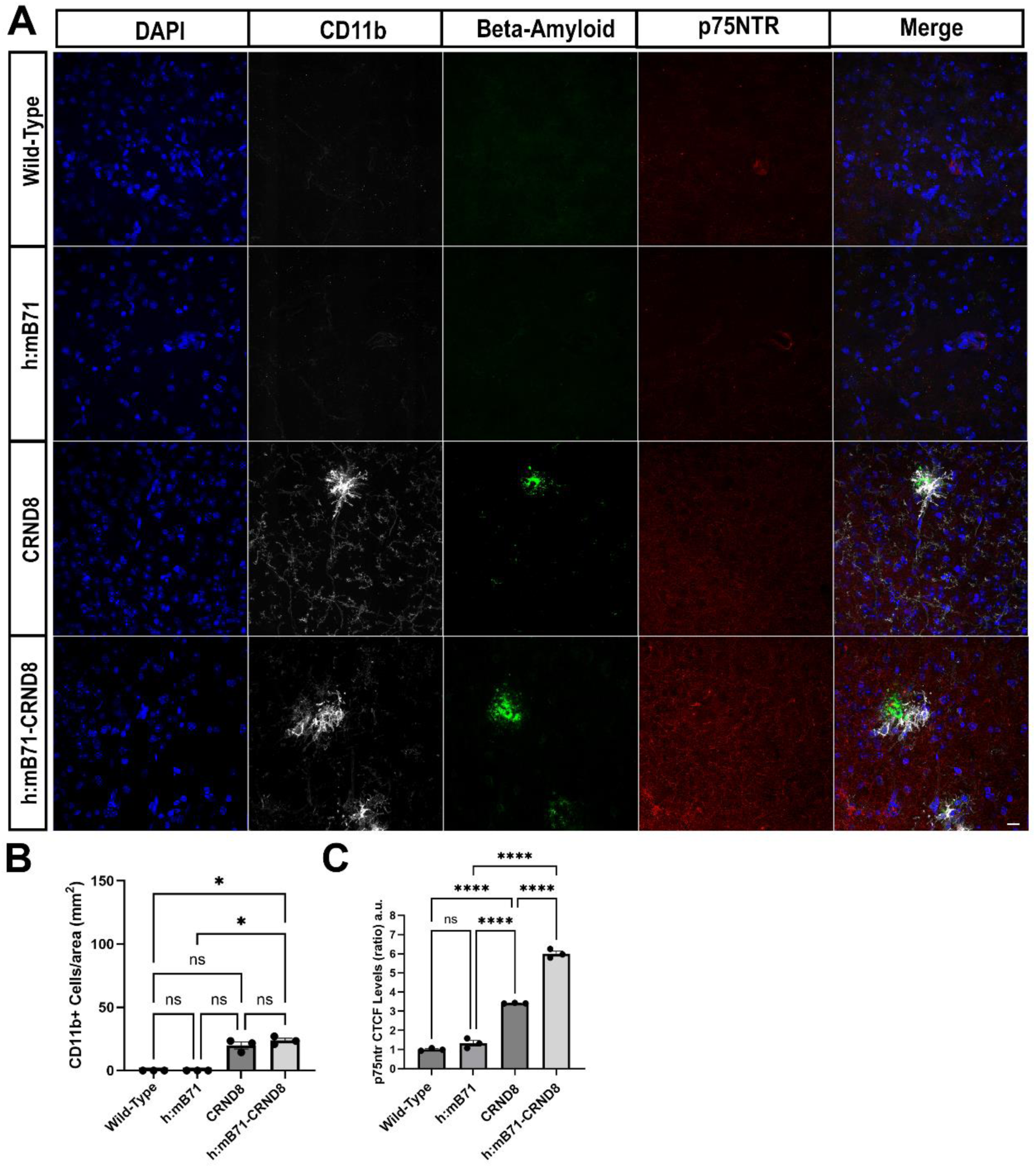
*At 3 months, h:mB71/CRND8 mice have more p75^NTR+^ compared to wt/CRND8, but comparably elevated CD11b+ cells in the dSubiculum*. (A) Representative confocal images of wt/wt, h:mB71/wt, wt/CRND8 and h:mB71/CRND8 mice showing DAPI stained nuclei, CD11b+ cells, Aβ deposition and p75^NTR^ upregulation. (B) wt/CRND8 and h:mB71/CRND8 express a significative upregulation of CD11b+ cells/area compares to wt/wt and h:mB71/wt mice, however, no difference was observed between h:mB71/CRND8 and wt/CRND8 mice. (C) In the dSubiculum of h:mB71/CRND8 mice more p75^NTR^ protein is detected as compared to wt/CRND8 mice. Data shown are mean +/− SEM. In (B)*p=0.0102, in (C) ****p<0.0001. One-way ANOVA followed by post hoc Tukey test. n.s not significant. N=3. Scale bars: 10µm

**Figure 6.**
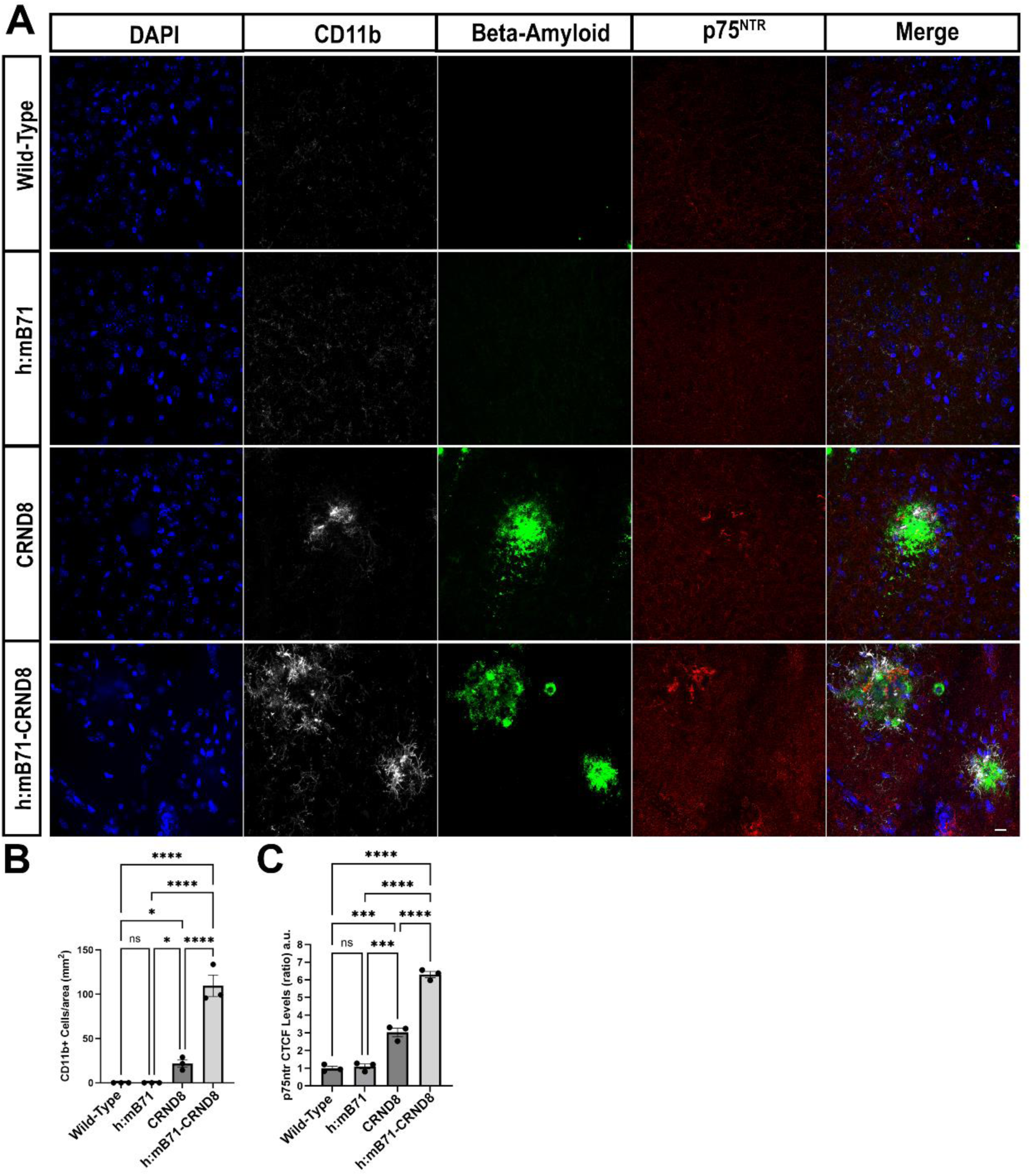
*h:mB71/CRND8 mice have more p75^NTR+^ and more CD11b+ cells in the dSubiculum compared to wt/CRND8 at 4mo*. (A) Representative confocal images of wt/wt, h:mB71/wt, wt/CRND8 and h:mB71/CRND8 mice showing DAPI stained nuclei, CD11b+ cells, Aβ deposition and p75^NTR^. (B) At 4mo, in the dSubiculum of the h:B71/CRND8 mice, there is significant upregulation of CD11b+ cells/area compared to wt/CRND8 (p=values), wt/wt and h:mB71/wt mice (p=values). Most of the CD11b+ cells are near beta amyloid accumulation. (C) In the dSubiculum of h:mB71/CRND8 mice there is more p75^NTR^ protein detected than in wt/CRND8 mice, however p75 is detected near beta amyloid accumulation in both genotypes. Data shown are mean +/−SEM. In (B)****p<0.0001, ***p=0.0246. In (C) ****p<0.0001; ***p=0.0003 (between h:mB71 and CRND8) and ***p=0.0002 (WT and CRND8). One-way ANOVA followed by post hoc Tukey test). N=3. Scale bars: 10µm

It is well established that inflammation occurs early during Aβ deposition in the CRND8 transgenic mouse model (Dudal et al 2004). To determine whether co-expression of h:mB7-1 altered neuroinflammation in either the onset or in the composition of activated cells, we assessed amyloid fibrils (beta amyloid), reactive astrocytes (GFAP) and microglia (Iba1), and activated microglia/infiltrating monocytes (CD11b) in the four genotypes of mice across 2-4 months of age. As expected, mice lacking the CRND8 transgene (whether wt/wt or h:mB7-1/wt) had no deposition of Aβ fibrils or plaques, and no abnormal accumulations of GFAP+ or Iba1+ or CD11b+ cells, at 2, 3 or 4 months of age, suggesting that the expression of h:mB7-1 in the absence of an inflammatory insult did not induce a glial or immune cell activation or recruitment (Fig. 2, 4, 5, 6 and Supplementary Fig. 2, 3, 4, 5). However, at 2 months of age, we found that the wt/CRND8 and h:mB7-1/CRND8 mice had modest increase in density of GFAP+ cells in the subiculum but no increases in Iba1+ cell (Supplementary Fig. 4, 5).

At 3 months of age, the amyloid plaque burden was readily detectable, and the density of GFAP+ cells in the h:mB7-1/CRND8 was markedly elevated (∼300% increase compared to wt mice) with a more modest induction of GFAP+ cell density in wt/CRND8 mice (∼150% increase compared to wt mice) suggesting that the expression of the h:mB7-1 allele enhanced astrocyte activation (Fig. 2 and 5). GFAP+ cells were localized both diffusely throughout the parenchyma, as well as frequently adjacent β-amyloid plaques. At this time point, CD11b+ cells were significantly elevated in both wt/CRND8 and h:mB7-1/CRND8 as compared to mice lacking the CRND8 transgene and detected adjacent to β-amyloid plaques (Fig. 5).

By 4 months of age, both wt/CRND8 mice and h:mB7-1/CRND8 mice continued to have elevated levels of GFAP (∼100% increase compared to mice lacking the CRND8 transgene) although there was no statistical difference between the two genotypes (Fig. 4). At this same time point, the h:mB7-1/CRND8 mice exhibited very high density of CD11b+ cells, exceeding the induction observed in wt/CRND8 mice by more than 4 fold (Fig. 6). Most of the CD11b+ cells in wt/CRND8 and h:mB7-1/CRND8 mice were adjacent to amyloid plaques. The expression of Iba1+ cells was also greater in the h:mb7-1/CRND8 mice as compared to wt/CRND8 (Fig. 7 and Supplementary Fig. 5C), suggesting that the expression of the h:mB7-1 allele augmented microglial activation at 4 months of age. Collectively these data suggest that in the setting of mutant Aβ (mice carrying the CRND8 transgene), the expression of h:mB7-1, as compared to murine B7-1, results in increased numbers of activated astrocytes at 2 and 3 months (Fig. 2, 4) followed by an increased numbers of activated microglia/macrophages at 4 months of age.

**Figure 7.**
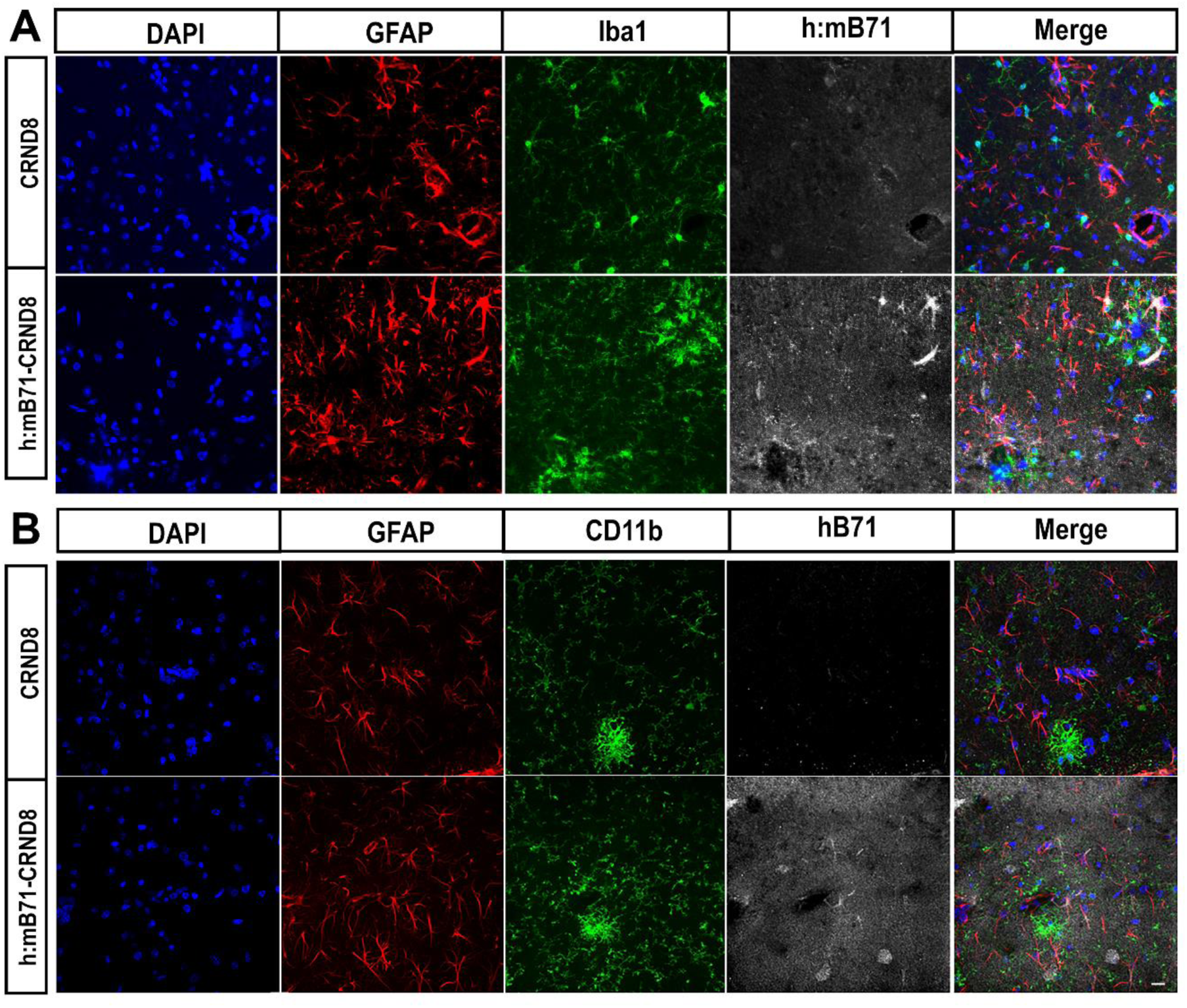
*hB71 protein is detected in GFAP+ cells and in GFAP- cells in the brain parenchyma at 4mo*. (A) Representative confocal images of wt/wt, h:mB71/wt, wt/CRND8 and h:mB71/CRND8 mice showing DAPI stained nuclei, Iba+ cells, GFAP+ cells and hB71 (B) Representative confocal images of wt/wt, h:mB71/wt, wt/CRND8 and h:mB71/CRND8 mice showing DAPI stained nuclei, CD11b+ cells, GFAP+ cells and hB71. N=3. Scale bars: 10µm.

### Expression of h:mB7-1 enhances synaptic elimination and dendritic remodeling in an AD model

Although no studies to date have focused on the subiculum, studies of hippocampal CA1 pyramidal neurons in the CRND8 mice demonstrate synaptic loss but no changes in dendritic length or arborization, as assessed in 3 month old male mice, a time point at which both contextual and cued memory is impaired (Steele et al 2014). Our prior studies using wt mice have demonstrated that direct injection of hB7-1 protein induces rapid elimination of apical dendritic spines of subicular pyramidal neurons, in a p75-dependant manner (Morano et al 2022). Thus, we sought to determine whether endogenous expression of h:mB7-1 results in a more pronounced reduction in spine number and dendritic morphology in the setting of mutant amyloid deposition and heightened neuroinflammation. To this end, Golgi analysis of the dorsal subiculum of mice from the 4 genotypes (wt/wt, h:mB7-1/wt, wt/CRND8 and h:mB7-1/CRND8) was undertaken. At 2 months of age, when h:mB7-1 expression was very low and the induction of neuroinflammatory markers was modest, the density and morphology of synapses in the apical dendrites of subicular pyramidal cells were comparable across genotypes (Supplementary Fig. 6). However, at 3 months of age, the h:mB7-1/CRND8 mice exhibited a 36% reduction of spine density as compared to wt/wt or h:mB7-1/wt mice, a value which was significantly greater than the 15% decrease in the wt/CRND8 mice, suggesting that expression of h:mB7-1 resulted in enhanced spine elimination (Fig. 8A and 8B). By 4 months of age, both h:mB7-1/CRND8 and wt/CRND8 mice exhibited comparable 40-45% reduction in spine density compared to wt/wt or h:mB7-1/wt mice (Fig. 9A and 9B). As activation of p75 on postsynaptic spines initially remodels spines from a mushroom to filopodial morphology prior to overt spine elimination (Giza et al 2018), we systematically characterized spines into three subtypes: mushroom, stubby or filopodial. A loss of mushroom spines was detectable in 3 mo old mice expressing the CRND8 transgene, and this effect was even more pronounced in h:mB7-1/CRND8 as compared to wt/CRND8 animals (Fig. 8C). By 4 months of age, both wt/CRND8 and h:mB7-1/CRND8 mice had comparable reductions in mushroom spines, and comparable modest increases in thin spines (Fig. 9C).

**Figure 8.**
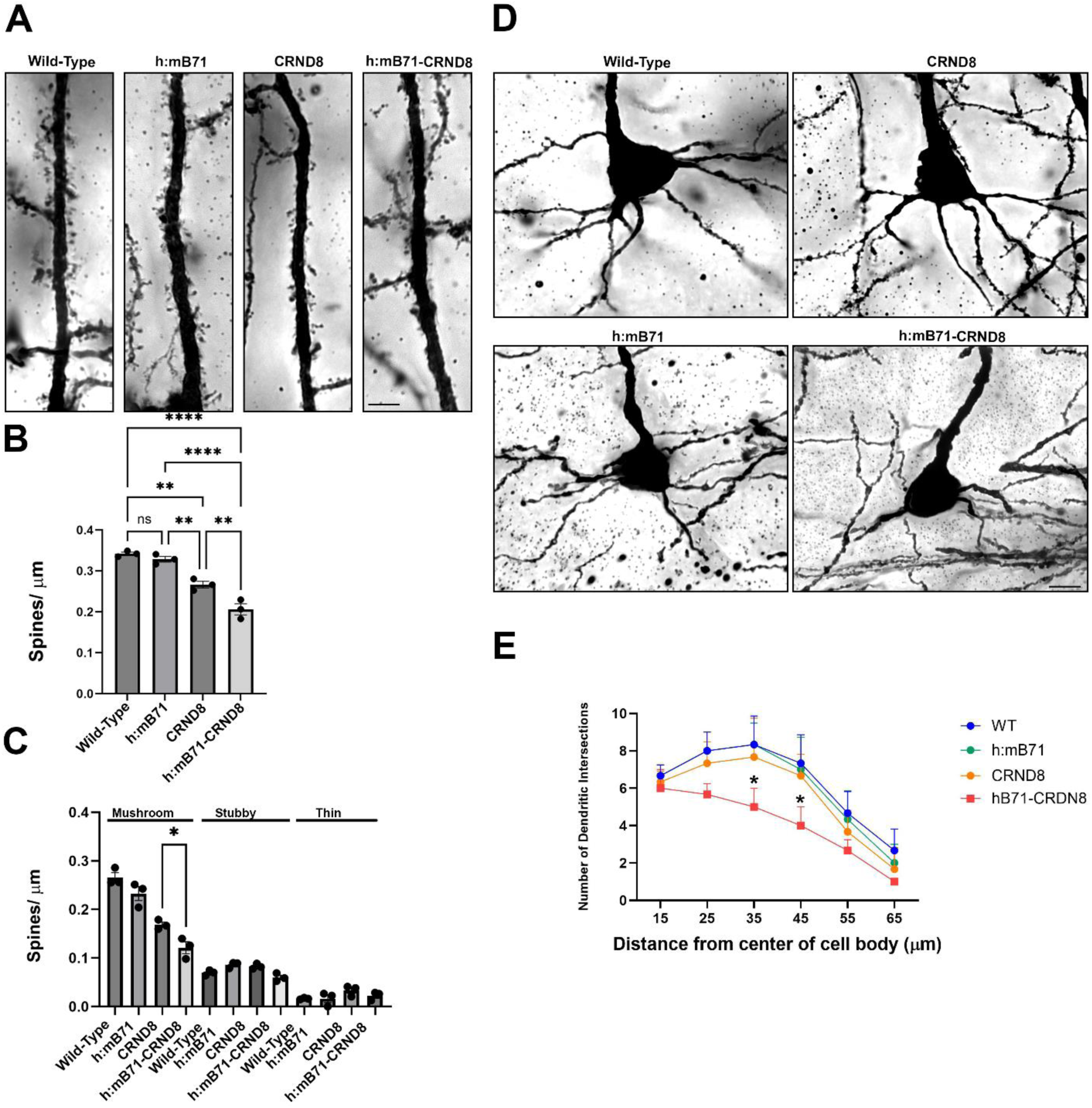
*h:mB71expression in the context of Aβ overexpression reduces total spine density and basal dendritic arborization in pyramidal subicular neurons.* (A) Representative Golgi-stained apical dendrites of neurons from wt/wt, h:mB71/wt, wt/CRND8 and h:mB71/CRND8 mice. (B) h:mB7-1/CRND8 mice exhibited a 40% reduction in spine density as compared to wt/wt or h:mB7.1/wt mice, and a 18% decrease compared to wt/CRND8 mice. (C) Quantification of mushroom, stubby or thin/filapodial spines. (D) Representative images of Golgi stained basal dendrites revealed reduced and altered branching patterns of from pyramidal subicular neurons in wt/wt, h:mB71/wt, wt/CRND8 and h:mB71/CRND8 littermates. (E) Quantification of the number of dendritic intersections. These results suggest that expression of h:mB7-1 in the context of neuroinflammation resulted in enhanced spine elimination and altered basal dendritic arbors. Data shown are mean +/− SEM. In (B)**p=0.0048 between h:mB71/wt and wt/CRND8; **p=0.0059 between wt/CRND8 and h:mB71/CRND8; ****p>0.0001. In (C) wt/CRND8 vs h:mB71/CRND8 *p=0.0441 (one-way ANOVA followed by post hoc Tukey’s test). In (E) *p=0.0274 showing comparison between wt/CRND8 and h:mB71/CRND8, 2 way ANOVA followed by post hoc Tukey’s test). N=3. Scale bars: 10µm.

**Figure 9.**
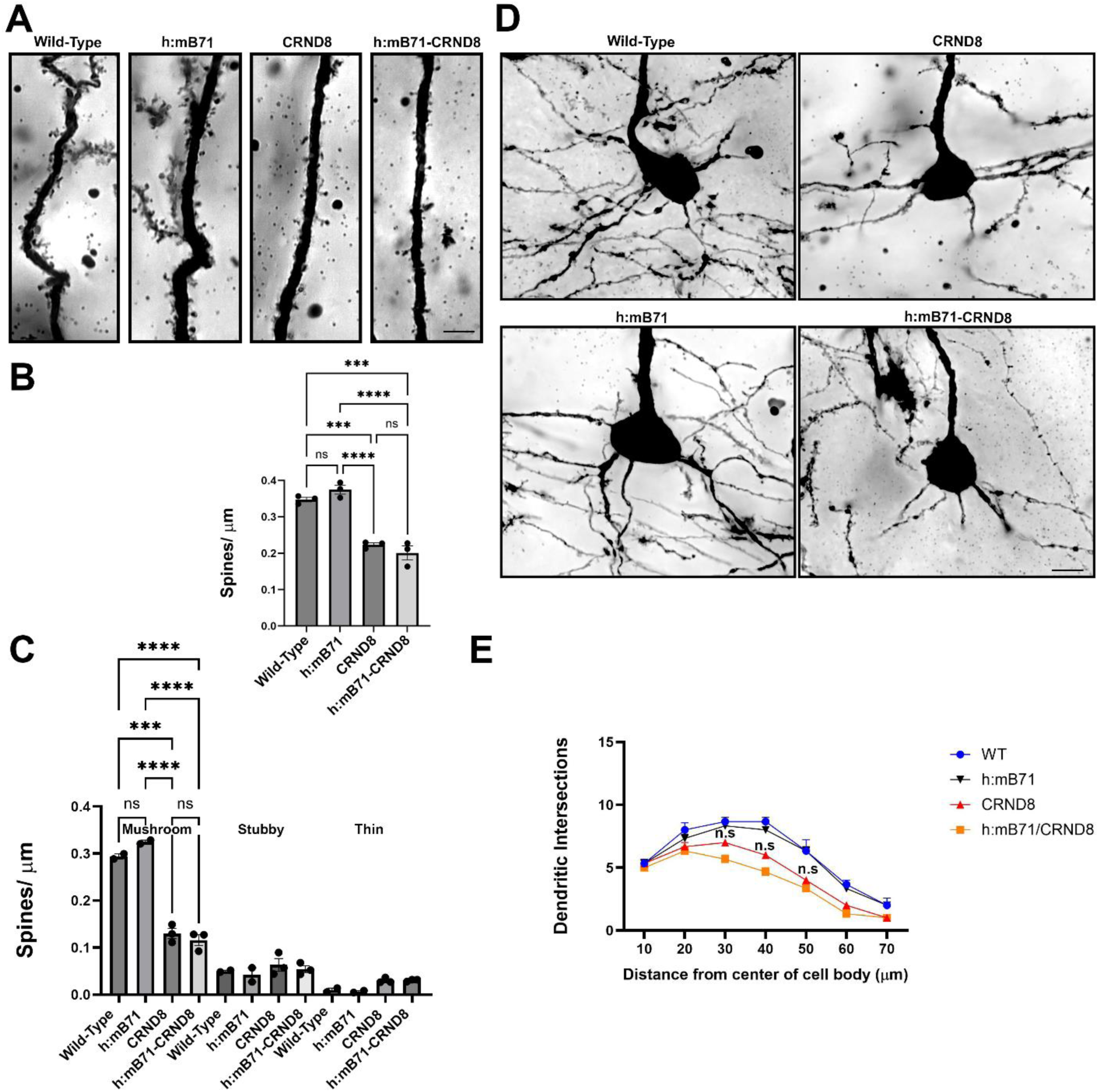
*At 4 mo mice expressing murine B7-1 or h:mB71 in the context of Aβ overexpression exhibit decreased total spine density and basal dendritic arborization in pyramidal subicular neurons*. (A) Representative Golgi-stained apical dendrites of neurons from wt/wt, h:mB71/wt, wt/CRND8 and h:mB71/CRND8 mice. (B) The h:mB7-1/CRND8 and the wt/CRND8 mice exhibited a 40% reduction of spine density as compared to wt/wtand h:mB7.1/wt mice. (D) Sholl analysis quantification revealed reduced and altered branching patterns of basal dendrites from pyramidal subicular neurons in wt/CRND8 and h:mB71/CRND8 compared to wt/wt, h:mB71/wt littermates. (E) Quantification of the number of dendritic intersections. No significant differences were found between wt/CRDN8 and h:mB71/CRND8 neurons along the dendritic arbor analyzed. Significant differences were found between wt/wt. h:mB71/wt and wt/CRND8 and h:mB71/CRND8. Data shown are mean +/− SEM. In (B)***p=0.004; ***p<0.0001, n.s not significant. In (C) ****p<0.0001. (one-way ANOVA followed by post hoc Tukey test). In (E) n.s not significant. N=3. Scale bars: 10µm

To determine whether h:mB7-1 expression had more extensive effects on neuronal morphology beyond synaptic elimination, we compared the complexity of dendritic branching of Golgi stained subicular pyramidal cells. Using Sholl analysis of basal dendritic arbors of pyramidal neurons in the dorsal subiculum at 15 to 70 microns from the cell body, neuronal arborization was comparable across genotypes at 2 months of age (Supplementary Fig. 6). However, at three months, h:mB7-1/CRND8 exhibited decreased complexity of arbors at 25-45 micron distances from the cell body, as compared to littermates wt/CRND8, wt/wt or h:mB7-1/wt mice (Fig. 8 D,E). At 4 months of age both h:mB7-1/CRND8 and wt/CRND8 mice exhibited decreased dendritic complexity from 25-60 microns from the cell body (Fig. 9 D,E). These results suggest that in addition to highly local effects on postsynaptic numbers and morphology, h:mB7-1 also has adverse effects on dendrite stability in the setting of amyloid deposition due to the mutant human transgene in CRND8 mice.

### Expression of h:mB7-1 promotes microvascular loss in an AD model

While historically the focus of studies has been on the functional impact of amyloid peptides on neuronal structure and function, these pathological peptides are also deposited in brain arterioles and capillaries of Alzheimer’s patients and in the CRND8 murine model (Chishti et al 2001, Farkas & Luiten 2001, Religa et al 2013, Zlokovic 2011). A limited analysis of vascular density in the cerebral cortices of CRND8 mice at 4.5 months of age documented a 30% reduction in endothelial cells, compared to wt mice (Religa et al 2013), suggesting that amyloid deposition, or the neuroinflammatory consequences thereof, impairs vascular architecture, with microvascular drop out. In addition, prior studies indicated that p75, normally absent in the adult vasculature, was induced following acute injury, specifically in models of ischemia/reperfusion injury of the heart and resulted in pericyte retraction and cardiac microvascular compromise (Siao et al 2012). At 3 months of age when Aβ deposition is established in mice expressing the CRND8 transgene, p75 and GFAP are detected in the region surrounding larger vessels in hippocampus, and h:mB7-1 + cells are localized to the perivascular region of h:mB7-1/CRND8 mice (Figs. 10, 11). To determine whether h:mB7-1 expression alters vascular structure, we undertook a quantitative analysis in the four genotypes of mice across 2-4 months of age. Image analysis of endothelial cells (using CD31 immunodetection) in the dorsal subiculum permitted an assessment of microvascular length and microvascular junctions (corresponding to capillary bifurcations). In 2-month-old mice, there were no differences in these microvascular parameters across genotypes (Supplementary Fig. 7). However, in animals of 3 months of age, a significant impact was observed in h:mB7-1/CRND8 mice compared to the other genotypes. Specifically, the capillary bed exhibited a 38% reduction in capillary length and a 43% decrease in capillary junctions, compared to animal lacking the CRND8 transgene (Fig. 12). No statistically significant differences were detected in capillary length or junctions in wt/CRND8 mice as compared to wt mice at 3 months of age. However, at 4 months of age we observed that the microvascular density of both the wt/CRND8 and the h:mB7-1/CRND8 mice were severely compromised to a similar extent, with a 40-45% reduction in vessel length and a 45-55% reduction in capillary junctions as compared to wt mice (Fig.13). To assess whether these deficits were restricted to endothelial cells, we evaluated the density and length of pericytes, using detection of PDGFRβ+ cells. Although no differences were observed across genotypes in 2 month old mice (Supplementary Fig. 7), h:mB7-1/CRND8 mice selectively demonstrated a significant loss of pericyte number (20%), and a reduction in pericyte length (30%) as compared to the other genotypes at 3 months of age (Fig. 14). These deficits were more pronounced in 4 months, with a more severe phenotype observed in h:mB7-1/CRND8 mice (70% reduction in pericyte number, and 60% reduction in pericyte length relative to wt mice) as compared to wt/CRND8 mice (40% reduction in pericyte number, 20% reduction in pericyte length relative to wt mice) (Fig. 14). Both endothelial cells and pericytes secrete extracellular matrix proteins, including laminin, as essential components of the basement membrane surrounding endothelial cells and pericytes, that function to maintain an intact blood brain barrier (Sebastiani et al 2015). Thus, we evaluated the density of laminin in direct contact with endothelial cells. While no differences were observed across genotypes in mice of 2 months of age (Supplementary Fig. 7), a profound decrease (65%) was observed in 3 month old h:mB7-1/CRND8 mice, but not in other genotypes (Fig.12). At 4 months, both wt/CRND8 and h:mB7-1/CRND8 mice demonstrated major reductions (70% wt/CRND8, and 80% h:mB7-1/CRND8 as compared to controls) in laminin detection. To directly assess impacts on microvascular perfusion, 3 month old mice of all four genotypes were injected intravascularly with anti-VE-cadherin, which binds to endothelial cells in patent vessels. While the wt/CRND8 mice exhibited a 33% reduction in VE-cadherin detection as compared to mice lacking the CRND8 transgene, the reduction in VE-cadherin staining was significantly greater in h:mB7-1/CRND8 mice (50% reduction compared to wt) (Fig. 12). Collectively, these findings suggest that animals expressing both the h:mB7-1 protein, and the CRND8 transgene exhibit an accelerated loss of microvascular integrity, impacting endothelial cells, pericytes and the basement membrane components of the neurovascular unit.

**Figure 10.**
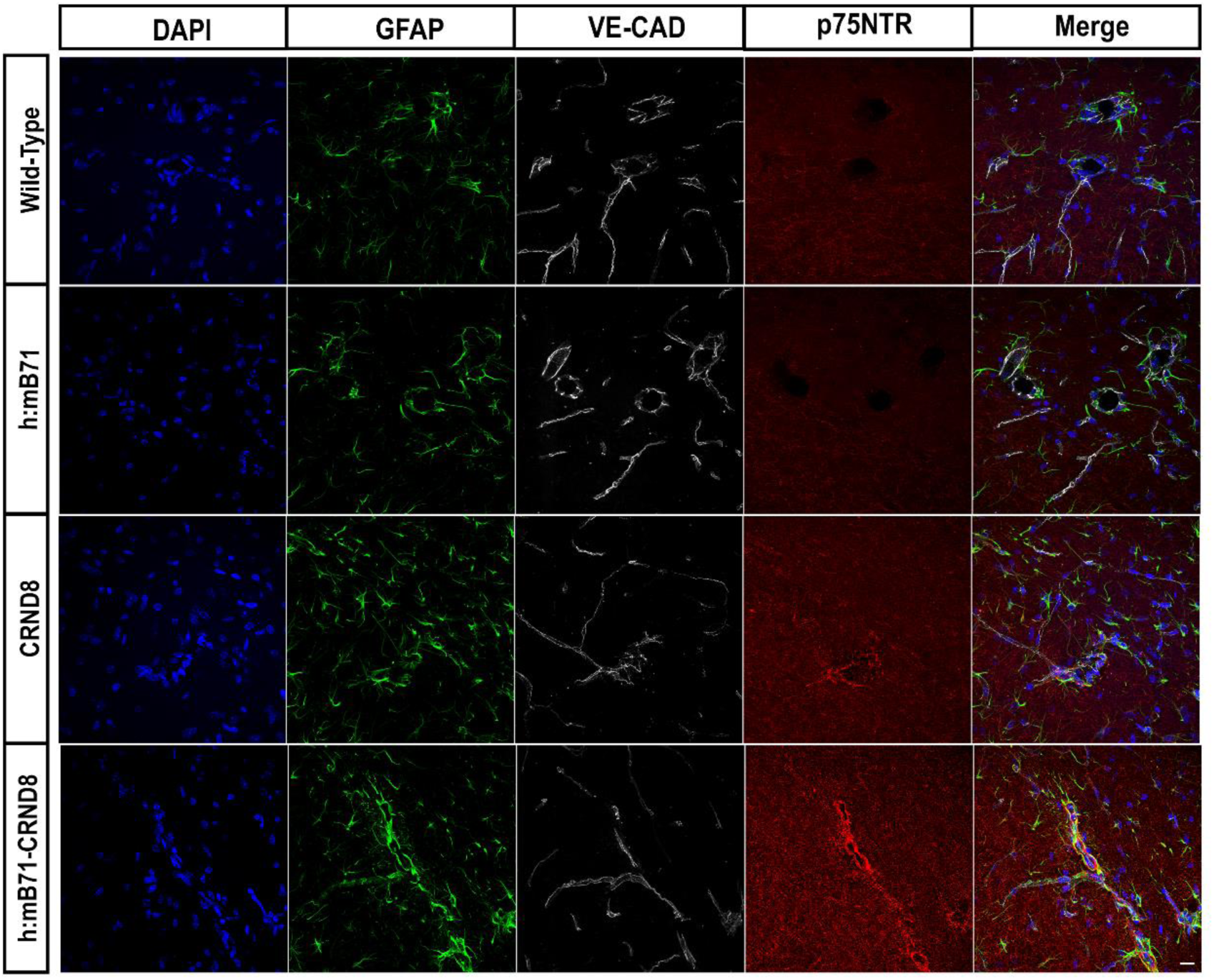
*In mice with Aβ overexpression, p75ntr protein is upregulated in the perivascular region of larger vessels in the hippocampus at 3mo*. Representative confocal images of wt/wt, h:mB71/wt, wt/CRND8 and h:mB71/CRND8 mice showing DAPI stained nuclei, GFAP+ cells, VE-CAD and p75^NTR^ protein. Scale bars: 10 µm

**Figure 11.**
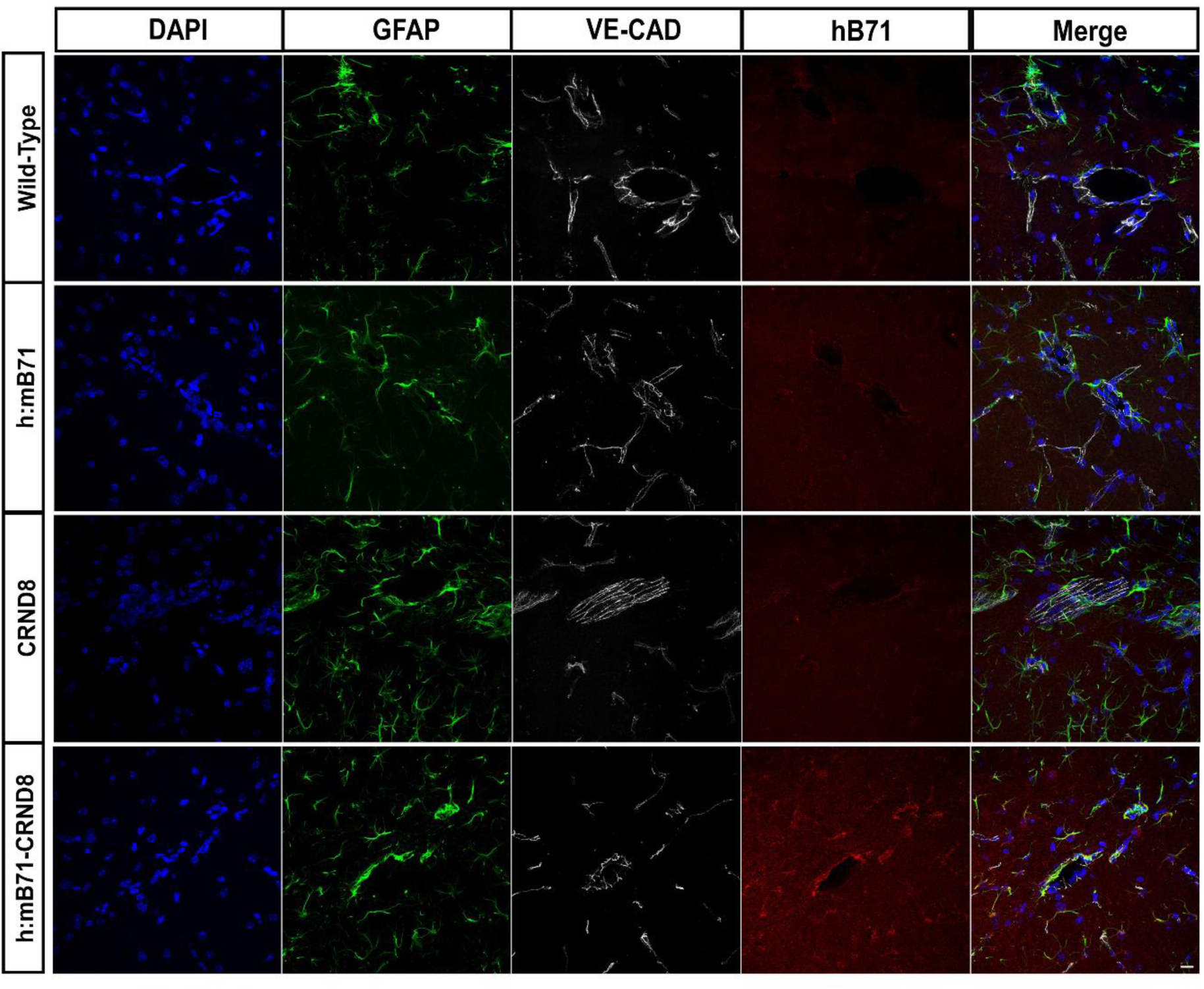
*In mice with Aβ overexpression h:mB7-1 protein is upregulated in the perivascular region of larger vessels in the hippocampus at 3 mo*. Representative confocal images of wt/wt, h:mB71/wt, wt/CRND8 and h:mB71/CRND8 mice showing DAPI stained nuclei, GFAP+ cells, VE-CAD and p75^NTR^ protein. Scale bars: 10 µm

**Figure 12.**
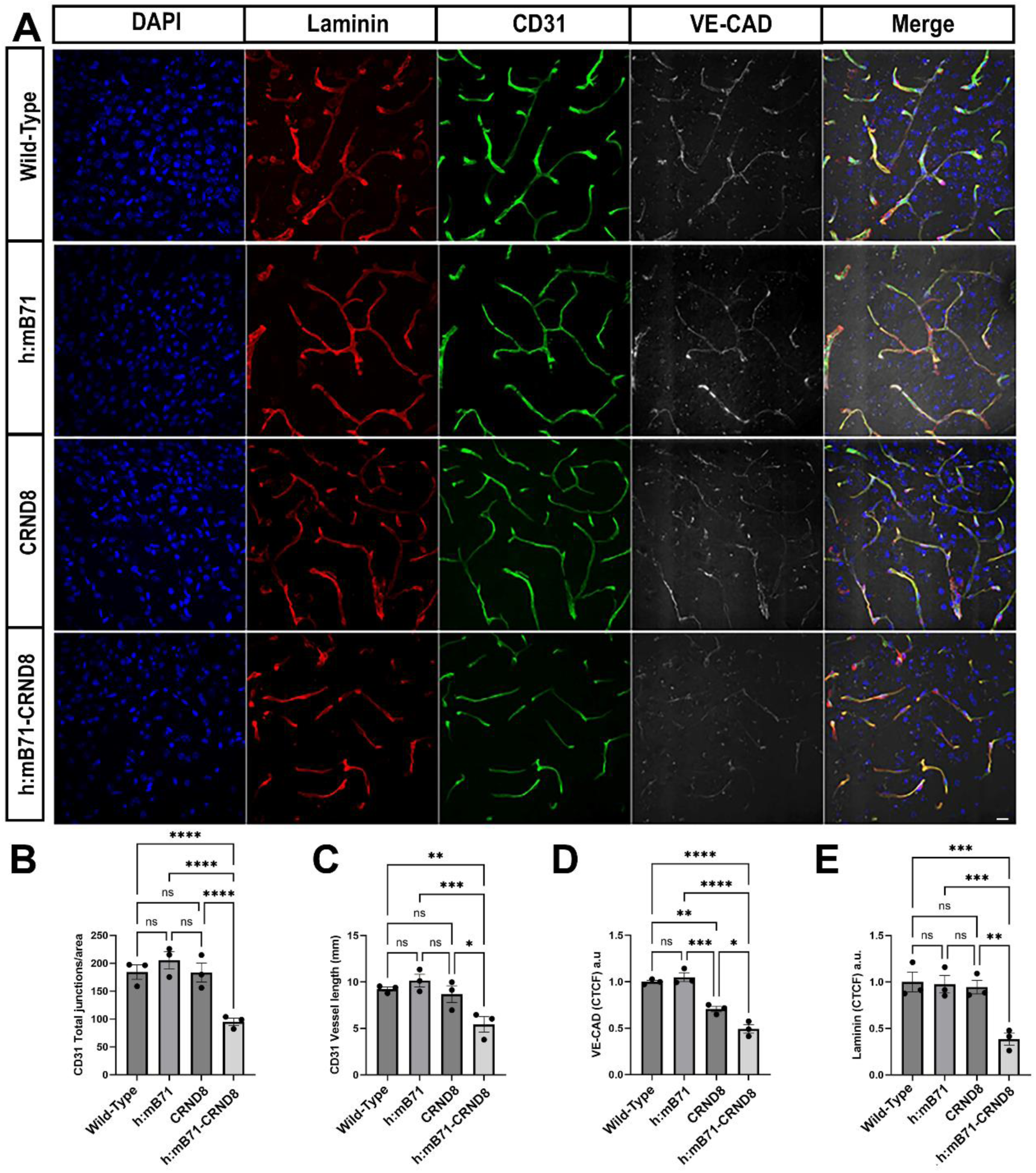
*Expression of h:mB7-1 promotes microvascular loss in the setting of Aβ overexpression in mice at 3mo*. (A) Representative confocal images of wt/wt, h:mB71/wt, wt/CRND8 and h:mB71/CRND8 mice showing DAPI stained nuclei, laminin, CD31, and VE-CAD expression. (B) and (C) h:mB71/CRND8 mice exhibit loss of microvascular junctions and length, as compared to wt/wt h:mB71/wt and wt/CRND8 mice. (D) There is significant loss of VE-CAD immunopositivity in h:mB71/CRND8 mice compared to wt/CRND8, wt/wt and hB71/wt mice, although wt.CRND8 mice exhibit a more modest reduction as compared to wt/wt and h:mB7-1 mice. (E) h:mB71/CRND8 exhibit loss of Laminin as compared to wt/wt, h:mB71/wt and wt/CRND8 mice. Data shown are mean +/− SEM. In (B)****p<0.0001. In (C) *p= 0.0144, **p=0.0027, ***p=0.0002. In (D) *p=0.0159, **p=0.0024, ***p=0.0009, ****p<0.0001. In (E) ***p=0.0003 (between wt/wt and h:mB71/CRND8), ***p=0.0005 (between h:mB7/wt and h:mB71/CRND8), **p=0.0011 (between wt/CRND8 and h:mB71/CRND8). One-way ANOVA followed by post hoc Tukey test. N=3. Scale bars: 10 µm.

**Figure 13.**
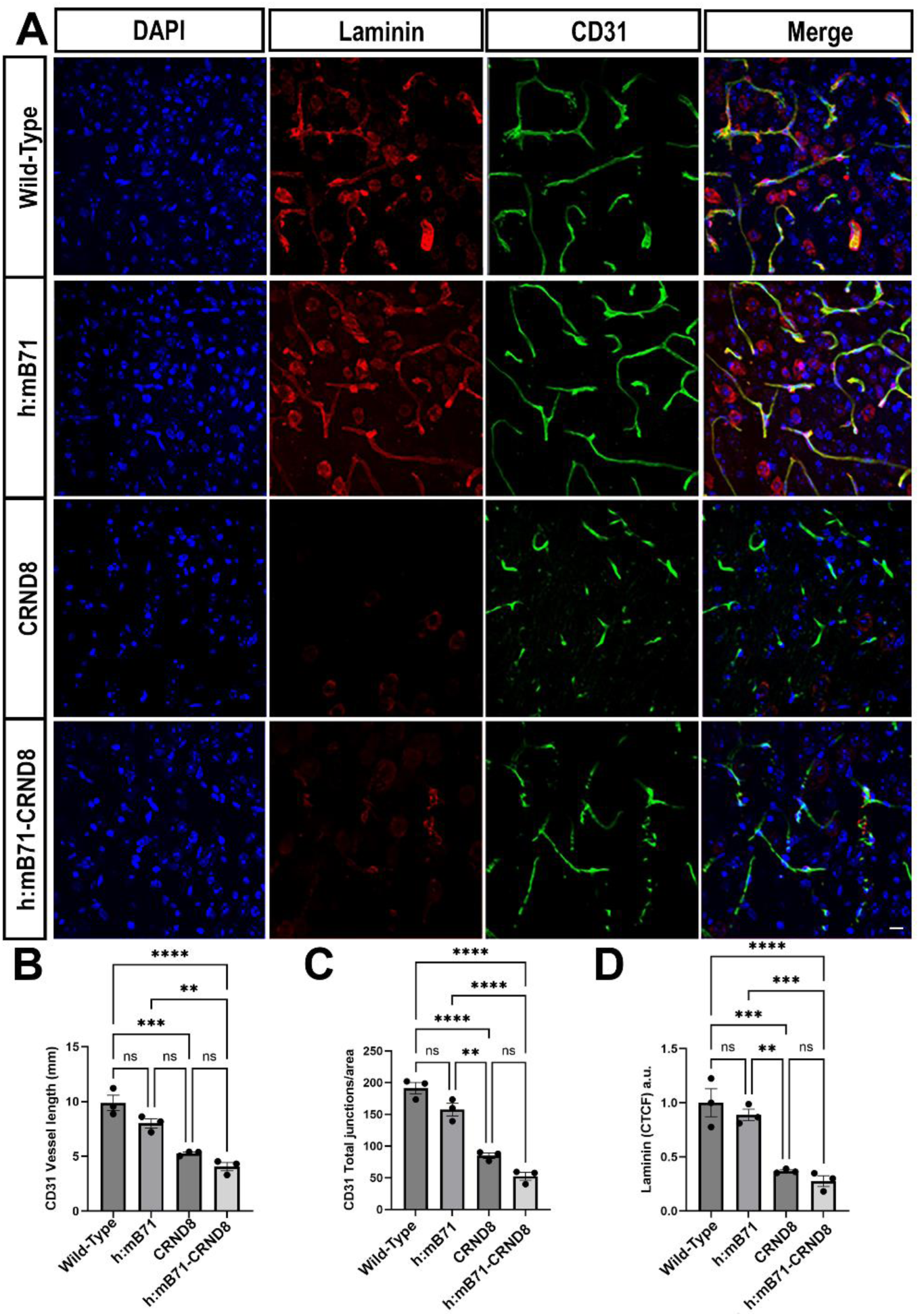
*h:mB7-1 promotes microvascular and laminin loss in mice with Aβ overexpression at 4mo*. (A) Representative confocal images of wt/wt, h:mB71/wt, wt/CRND8 and h:mB71/CRND8 mice showing DAPI stained nuclei, laminin and CD31 expression. Upon expression of the CRND8 transgene, there is a significant loss of vessel length (B), junctions (C) and laminin (D) compared to wt/wt and h:mB71/wt mice. No significant differences were found between h:mB71/CRND8 and wt/CRND8. Data shown are mean +/− SEM. In (B)**p 0.0018, ***p=0.0002, ****p<0.0001. In (C) ****p<0.0001, **p=0.0012. In (D) **p=0.0027, ***p=0.0002 (between wt/wt and wt/CRND8), ***p=0.0003 (between h:mB71/wt and h:mB71/CRND8), ****p<0.0001. One-way ANOVA followed by post hoc Tukey test. N=3. Scale bars: 10 µm.

**Figure 14.**
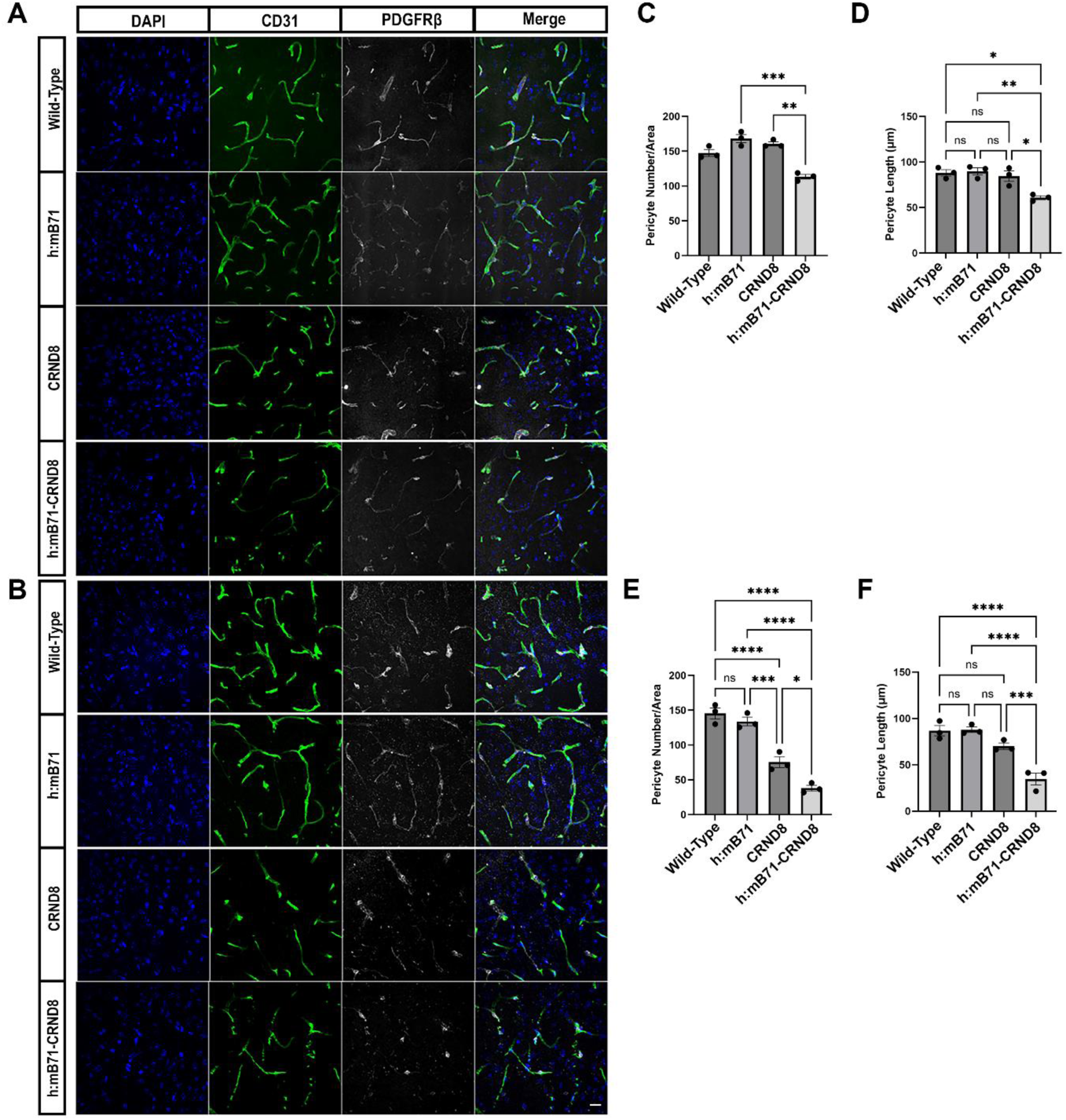
*h:mB7-1 promotes loss of pericyte length and number in mice with Aβ overexpression at 3 months and 4 months of age.* (A) and (B) Representative confocal images of wt/wt, h:mB71/wt, wt/CRND8 and h:mB71/CRND8 mice showing DAPI stained nuclei, CD31 and PDGFRβ (pericyte) expression. (C) At 3 mo, h:mB71/CRND8 mice exhibit loss of pericyte number/area and (D) pericyte length compared to wt/CRND8 (and to wt/wt and h:mB71/wt mice). At 4mo, the h:mB71/CRND8 mice exhibit a significant decrease of (E) pericyte number/area and (F) pericyte length compared to wt/CRND8, wt/wt and h:mB71/wt mice. Data shown are mean +/−SEM. In (C)**p=0.0035, ***p=0.0005. In (D)*p=0.0147 (between wt/wt and h:mB71/CRND8), *p=0.0494 (between wt/CRND8 and h:mB71/CRND8) **p=0.0081. In (E) *p=0.0352, ***p=0.002, ****p<0.0001. In (F) ***p=0.0007, ****p<0.0001. One-way ANOVA followed by post hoc Tukey test. N=3. Scale bars: 10 µm.

## Discussion

The expression of a human:mouse B7-1 chimeric protein which is capable of binding to the p75 receptor leads to a phenotype of accelerated neuroinflammation when co-expressed in a well characterized murine model of mutant Aβ overexpression. These findings suggest that this primate-restricted interaction of B7-1:p75 may be clinically relevant and could be important in mechanistically dissecting mediators of inflammation which occur in the human brain, but have not, to date, been accurately represented in murine models. As such, our findings suggest that the human B7-1:p75 interaction is a new therapeutic target, that accelerates neurodegeneration, enhances activation of resident glial, and leads to microvascular compromise in the subiculum, a region which is involved in learning and memory and significantly affected in early stages of AD.

Synaptic pathology is a hallmark of prodromal and overt AD. Oligomeric Aβ is toxic and can interact with multiple post-synaptic proteins including mGluR5 and NMDA receptors (Benarroch 2018, Spires-Jones & Hyman 2014). However, it is well established that the p75 receptor is expressed on post-synaptic spines in the hippocampus (Giza et al 2018). While prior reports suggest that p75 can be activated by high concentrations of Aβ (Patnaik et al 2020, Perini et al 2002),our findings suggest that p75 may play an additional prominent role in accelerating synaptic loss in humans utilizing locally expressed human B7-1. The effects of Aβ oligomers are well-established on the intrinsic properties on CA1 pyramidal neurons of CRND8 mice in the prodromal phase (2.5 months), and on long-term potentiation in overt disease (5 months) (Jolas et al 2002, Wykes et al 2012). It will be important to determine in future studies whether there are specific effects on synaptic plasticity due to local h:mB7-1 expression and activation of post-synaptic p75, as has been established with other p75 ligands such as proBDNF, which induces long term depression (Woo et al 2005, Yang et al 2014). As hB7-1 may remain cell tethered, in contrast to a diffusible ligand such as proBDNF, its effects on neuronal morphology may predominate, as we have observed in the h:mB7-1/CRND8 mouse model which exhibits not only synaptic loss, but also accelerated negative effects on dendritic arborization, with overt loss of dendrites in the subiculum (Fig. 8 and 9).

It is well recognized that the neuroinflammatory process in AD results from the dysregulation of multiple cell types, including astrocytes, microglia and peripheral immune cells of hematopoietic origin which invade the brain parenchyma (Heneka et al 2025, Vara-Perez & Movahedi 2025, Yan et al 2022). Despite investigation of the role of immune cells in murine models of AD (Chittimalli et al 2024), it is clear that mouse models only partially recapitulate the complex brain environment encountered in patients, which may be due in part to the limited homology of immune ligands across species, as is the case for B7-1 (Ernst & Carvunis 2018, Morano et al 2022). However, the expression of h:mB7-1 results in an accelerated pattern of activation in multiple cell types, rather than being restricted to only a single class of resident glial cells. This suggests that hB7-1/p75 activation downstream of a singular inflammatory insult (expression of high levels of amyloidogenic Aβ) may augment inflammatory signaling in multiple cells, including microglia and astrocytes. The temporal pattern of hB7-1-induced enhanced activation, affecting GFAP+ cells at 3 months and CD11b+ cells at 4 months suggests that numerous classes of cells may differentially regulate B7-1 or p75. Further investigation will be required to determine which neuroinflammatory effects are direct (via hB7-1:p75 engagement) or utilize local induction and release of cytokines to activate numerous types of glia. The consequences, however, of enhanced activation are clear, with increased synaptic pruning and dendrite loss, and microvascular compromise. Furthermore, the ability of hB7-1 to accelerate neuroinflammation in what would be considered a prodromal phase in this mouse model suggests that the hB7-1/p75 interaction could be an important therapeutic target.

The accelerated microvascular impairment observed in the h:mB7-1/CRND8 mice as compared to the wt/CRND8 is particularly relevant as AD is accompanied by a reduction in microvascular perfusion in association with white matter attenuation and impaired cognition. With significant recent advances in imaging of patients with prodromal Alzheimer’s disease, it is now recognized that elevated amyloid β levels in patients correlate with decreased microvascular blood flow and increased heterogeneity of flow in distinct brain regions (Madsen et al 2023) underscoring impact of microvascular dysfunction in this disease. The maintenance of microvascular flow requires an intact neurovascular unit, the product of numerous cell types which can be impacted by Aβ levels. This vascular dysfunction is correlated with (a) amyloid β accumulation in the wall of arterioles and capillaries as cerebral Aβ angiopathy (Kisler et al 2017); (b) contraction of brain pericytes leading to a reduction in pericyte coverage (Nortley et al 2019, Uemura et al 2020);(c) injury to the basement membrane of capillaries, leading to a dysfunctional blood brain barrier (Zenaro, Piacentino & Constantin 2017) as well as infiltration of peripheral immune cells to the brain parenchyma (Cao & Zheng 2018); (d) attenuation and withdrawal of astrocyte foot processes, leading to impaired metabolic function of neurons (Lee et al 2023, Ziar, Tesar & Clayton 2025); and (e) microglial activation leading to pericyte dysfunction (Morris et al 2023). The expression of h:mB7-1 may play a direct role in several of these processes as we observe accelerated loss of basement membrane components, enhanced activation of astrocytes and microglia, and accelerated loss of pericytes in h:mB7-1/CRND8 animals as compared to wt/CRND8 mice. The mechanistic dissection of this will be assisted by more detailed localization of both ligand and receptor, although published ssRNA seq data from the brains of AD patients data suggests that activated microglia are a prominent source B7-1 (CD80), and perivascular fibroblasts as well as other vascular cells express p75 (Yang et al 2022).

Our study has several limitations which may result in an underestimate of the potential neuroinflammatory effects of hB7-1:p75 engagement in humans. First, the breeding strategy used to generate the four cohorts of mice resulted in h:mB7-1/CRND8 mice that are heterozygous for h:mB71, with one allele that encodes murine B7-1 which is incapable to binding to and activating p75. Thus, this murine model is likely to underestimate the role of B7-1 in humans, expressing two B7-1 alleles. Second, the premature lethality of h:mB7-1/CRND8 mice (only 25% survival at 4 months), suggests that there may be more extensive impacts in the brain, which could not be assessed in the mice surviving to the 3 and 4 month time points. It is possible that there are additional pathological conditions which will require a more extensive future analysis in extra-neuronal tissues. These additional studies may strengthen our understanding of the role of human B7-1 in the setting of aberrant Aβ expression in Alzheimer’s disease.

## Methods

### Generation of a h:mB7-1 chimeric mouse

The region of B7-1 which binds to p75 is the IgV domain, and the murine IgV domain is encoded by murine exon 2 (106 residues), and the human IgV domain is encoded by human exon 3 (105 residues) (**Fig. 1B**). The B7-1 human-exon3-replace-mouse-exon2 allele was generated using CRISPR/Cas9 mediated genome editing ^[1]^ by the Einstein Gene Targeting Facility. A guide RNA (gRNA) targeting intron 1 of mouse B7-1 gene, named B7-1 gRNA 85-58 (targeting sequence: CACTATAAAGTGACTAACCACGG) and a gRNA targeting intron 2 of mouse B7-1 gene, named B7-1 gRNA 75-56 (targeting sequence: AACGTCTGACTTCATAAGTGAGG respectively, were designed by an online software Benchline (www.benchline.com) and were generated by in vitro transcription. A single stranded donor DNA that contains around 800 nt homologous arms and the surrounded Human B7-1 exon 3, named B7-1 donor DNA, was generated by SLiCE and ivTRT. Cas9 protein was purchased from PNB. All the above CRISPR ingredients were mixed and injected into the fertilized eggs of C57BL/6J mice, and then the injected fertilized eggs were transferred to the pseudopregnant CD1 female mice for producing pups. The resulting progeny were screened by DNA sequencing to identify correctly targeted founder (F0) mice. Three founder lines were obtained, and one was discarded following sequencing as it encoded a mutation. The remaining two founders were evaluated by Southern blot (Fig. 1C) to confirm that inadvertent mutagenesis had not occurred. Founders have been maintained on C57BL/6J for more than 9 generations. Genotyping of mice is undertaken using PCR: PCR with primers B7-1 GTF (recognize the upstream flanking region of B7-1 donor DNA in <u>mouse</u> B7-1 intron 1) and Hb7-1 KIR (recognize the <u>human</u> B7-1 exon 3) to yield a targeted allele of 578 bp using primers B7-1 GTF (ttgagacagagcctcccagt) and Hb7-1 KIR(tgccagtagatgcgagtttg).CRND8 mice were obtained from the University of Toronto, and maintained on a CEH-He/C57Bl6 background.

### Study approval

All of the animal experiments were conducted in accordance with the Guide for the Care and Use of Laboratory Animals (National Academic Press, 2011) and approved by the Institutional Animal Care and Use Committee at Weill Cornell Medical College.

### Titrations with recombinant CTLA-4, CD28, and p75^NTR^ protein

HEK-293 Freestyle cells (Invitrogen) were transiently transfected with constructs for human B7-1 GFP, mouse B7-1 GFP or h:m chimera B7-1 GFP. Two days post transfection, cells were centrifuged (500g, 5min) and resuspended to 2×10^6^ cells/mL in 1X PBS with 0.2% BSA. Recombinant mCTLA-4-hIgG1, mCD28-hIgG1, and mp75^NTR^-hIgG1 (Sinobiological) were added at the indicated concentrations to 100,000 transfected cells (50 µL) plated in 96-well V-bottomed plates. Protein receptors were bound to B7-1 expressing cells for 1 hour at 4°C with shaking at 900 rpm. Proteins were then removed by centrifugation and anti-human Alexa 647 antibody (1/200 dilution, Invitrogen) was added and bound for 30 min. After binding, cells were washed with 1X PBS, 0.2% BSA three times and immediately analyzed by flow cytometry. The percent bound (percentage of GFP positive cells that were also Alexa 647 positive) was normalized and used to determine EC50 values.

### Flow cytometry

Spleens were harvested from lethally anesthetized mice, and splenocytes from WT and h:mB7-1 animals were isolated and activated with either 1x lipopolysaccaharide (LPS) or a combination of 10 ng/mL phorbol 12-myristate 13-acetate (PMA) and 250 ng/mL ionomycin. Cells were activated for 24-hours, collected and stained with antibodies against 1) mCD45-AF647 (all differentiated hematopoietic cells), 2) mCD3-APC/Cy7 (all T-lymphocytes), 3) mCD19-AF488 (all B-lymphocytes), 4) CD11b-AF700 and F4/80-BV650 (macrophages), 5) CD11c-BV570 (dendritic cells), and 6) either anti-mB7-1 or hB7-1- PE Dazzle. After staining with antibodies for 30min at 4°C, cells were pelleted and stained with LIVE/DEAD Violet (Invitrogen) for 30min, washed three times with 1X PBS, 0.5%FBS, 1mM EDTA and analyzed by flow cytometry. Cells were gated by forward and side scatter, then “Live”, then singlets (FSC-H v FSC-A, and SSC-H v SSC-A) and then all CD45 positive. Cells were then subsequently gated into the following populations: T-cells (CD3+, CD19-), B-Cells (CD3-, CD19+), macrophages (CD3-, CD19-, CD11b+, F4/80+), and dendritic cells (CD3-, CD19-, CD11b-, F4/80-, CD11c+). B7-1 expression was then gated for each population and the geometric mean of the PE Dazzle florescence (the amount of B7-1 antibody bound) was normalized to the untreated cell populations. Data shows the average from 2 technical replicates from 7 wild-type and 8 h:mB7-1 chimera mice.

### Antibodies

The following primary antibodies were used: beta amyloid polyclonal antibody (CT695) rabbit Invitrogen ref. 51-2700, 1.0 μg/mL o.n., CD31 (R&D Systems, AF3628) - 1 µg/mL, overnight (o.n.) incubation, hB7-1 (R&D Systems, AF140)-7.5 mg/ml, 2 day incubation (followed with secondary biotinylated and streptavidin conjugated Alexa Fluor®), Iba1 (NBP2-19019)-1:1000, Laminin (NB300-144)- 1:250, o.n. incubation, mouse B7-1 (R&D Systems, AF740)- 2.5 mg/mL, o.n, anti-p75 (R&D Systems, AF1157)- 1 ug/mL, o.n, anti-PDGFRb (ThermoFisher, 14-1402-82)- 1:200, 2 day incubation, GFAP (Sigma, G3893)- 1:400, o.n., CD11b (Biolegend, 163702) (1:500). All the primary antibodies were incubated at 4^0^C, PBS, 0.08% Triton, 1%BSA.

The following secondary antibodies were used: Invitrogen Alexa Fluor® 647 donkey anti-rat IgG (H+L) ref. A78947, Invitrogen Alexa Fluor® 568 donkey anti-rabbit IgG (H+L) ref A10042. Invitrogen Alexa Fluor® 647 donkey anti-goat IgG (H+L) ref A21447, Invitrogen Alexa Fluor® 594 donkey anti-mouse IgG (H+L) ref. A21203, Donkey anti-Goat IgG (H+L) Cross-Adsorbed Secondary Antibody, Biotin Invitrogen (A16009) 1:1000; Streptavidin Alexa Fluor® 647 (S21374) or 594 (S11227), Invitrogen Alexa Fluor® 568 Donkey anti-mouse IgG (H+L) ref A10680. All the secondary antibodies were incubated at RT, in 1% BSA/PBS.

### Trans-cardiac perfusion and tissue sectioning

Mice were deeply anesthetized with sodium pentobarbital and perfused transcardially using the Perfusion Two Automated Pressure Perfusion system (Leica Microsystems) with ice-Cold PBS, followed by of 4% paraformaldehyde (pH 7.4). The brains were then removed and post-fixed with 4% paraformaldehyde at 4°C overnight and transferred to a sucrose solution (30% sucrose in PBS) at 4°C for 48 to 72 hours. Coronal sections (thickness, 30 μm) were prepared using a freezing microtome. The sections were set in an antifreeze solution (30% glycerol, 30% ethylene glycol, and 40% 0.25M PB) and stored at –20°C. Free-floating serial sections were washed (3 times for 10 minutes each) in PBS and incubated for 1 hour in a blocking solution containing 1% BSA in PBS with 0.08% Triton X-100 (PBS–Tx), following incubation in primary antibodies diluted in the blocking solution mentioned above and incubated 24 to 48 hours at 4°C with rotating. After washing 3×10 min in PBS (rotating), sections were incubated for 45mi-1 hour with secondary antibodies and then washed 3×10 min in PBS (rotating). The sections were mounted and air-dried overnight in the dark. Slides were coverslipped using water-soluble glycerol-based mounting medium containing DAPI and sealed with nail polish.

### Retro-orbital injection of anti-VE-CAD

The mice were anesthetized with isoflurane and anti-VE-CAD (Biolegend Alexa Fluor 647 anti mouse (VE-cadherin) clone BV13 Isotype rat IgG cat. 138006) - 1:5 dilution in PBS, was retro-orbitally injected using a tuberculin syringe with a 27-gauge. Mice were monitored as they recovered for 20min before deeply anesthetizing and performing trans-cardiac perfusion with PBS and 4%PFA as described above.

### Brightfield and fluorescence microscopy

Golgi-stained sections were imaged using Ni-E Microscope, with Camera Nikon DS-Qi2, using Plan Fluor 100x Oil DIC H. Acquired images were processed and reconstructed using FIJI(ImageJ) and Photoshop Align and merge functions. Immuno-stained sections were image using Ti2 Microscope spinning disk confocal, with a Plan APO λD 20x OFN25 DIC N2 and Plan Apo λD 60x Oil OFN25 DIC N2. The laser lines used were 405 nm, 488 nm, 561 nm, and 639nm. The images were processed and quantified using Fiji (ImageJ) scientific software.

### Immunofluorescence and Microvasculature analysis

The CD31 stained images were analyzed with AngioTool software. CD11b+ cells, GFAP+ cells, and pericytes number quantification were done using FIJI(ImageJ) using Analyze particles function; Pericytes length quantification were done using FIJI(ImageJ) manually measuring the length of the pericyte, with the line measuring tool; p75, Laminin and VE-CAD protein levels were quantified using FIJI(ImageJ) to calculate CTCF = Integrated Density – (Area of selected cell X Mean fluorescence of background readings). All the data were normalized to the area quantified.

### Golgi analysis

Neurons were labeled using the Golgi-Cox method as previously described (Chen et al, 2006). Golgi impregnated sections were numbered in a blinded manner prior to quantitative analysis. Neurons for dendritic arborization analysis were selected if there was a cell body with primary dendrites emerging from the soma without transection of the dendrites, there was little overlap with neighboring neurons, and impregnation was consistent along the dendrites. 20-30 neurons from each animal were acquired and the images were reconstructed using FIJI (imageJ) scientific software and Photoshop (using Align, Blend and Merge functions) Using sections prepared for Golgi impregnation, dendritic spines of pyramidal cells in the dorsal subiculum were counted at 100X magnification using a Golgi-stained sections were imaged using Ni-E Microscope, with Camera Nikon DS-Qi2, using Plan Fluor 100x Oil DIC H. Isolated dendrites were selected and the total number of dendritic spines along a 100 micrometer length was counted. The Sholl analysis was conducted on basal dendritic arbor using FIJI (imageJ) scientific software, Neuroanatomy Shortcuts Tool, Sholl Analysis functions.

### Quantification and statistical analysis

Sample sizes (n) indicated in figure legends correspond to the number of mice used in the experiments. All statistical parameters are presented as means ±SEM (Standard Error of Mean). All data were analyzed with GraphPad Prism 10.6.1 software (San Diego, CA, USA). Statistical significances were calculated via One Way ANOVA and 2-Way ANOVA, with Tukey’s post hoc tests to control for multiple comparisons (for three or more group comparisons), or two-way ANOVA with Tukey’s post hoc tests (to assess statistical significance between means), as indicated within individual figure legends. In figures, asterisks denote statistical significance marked by *, **, ***, **** (p values in figures) and “n.s.” indicates no statistical significance.

## Acknowledgements

We thank Alison Looke for assistance in maintaining mice, Lino Tessarollo and Mary Ellen Palko for experimental discussion and Southern blot analysis and Theresa Milner for critical discussions of brain ultrastructure. We gratefully acknowledge Yongwei Zhang and Winfried Edelmann in the Albert Einstein Gene Targeting Facility for assistance in the generation of the hm:B7-1 chimeric knock-in mouse. This work was supported by the NIH grants 1RF16078613 (to BLH, FSL and SCA and Einstein Macromolecular Therapeutics Development Facility.

## Author contributions

VD, SCGT, SCA, FSL and BLH conceived the project and designed the experiments. VD performed the immunofluorescence studies, assisted by PK, RSS, RC, and VD, BLH, FSL, SCA, SCGT reviewed this research. NCM and SCGT performed the flow cytometry analysis using cell lines and primary splenocytes. BLH, SCA, FSL and NCM assisted in designing the strategy for the chimeric mouse. RSS, RC, VD conducted the mouse breeding and survival analysis and VD developed and oversaw the breeding strategy. FSL, BLH and SCA provided funding, and FSL, BLH, SCA, SCGT and VD generated and revised the manuscript. All authors approved the final manuscript.

## References

Andrews SJ, Renton AE, Fulton-Howard B, Podlesny-Drabiniok A, Marcora E, Goate AM. 2023. The complex genetic architecture of Alzheimer’s disease: novel insights and future directions. EBioMedicine 90: 104511

Bellucci A, Luccarini I, Scali C, Prosperi C, Giovannini MG, Pepeu G, Casamenti F. 2006. Cholinergic dysfunction, neuronal damage and axonal loss in TgCRND8 mice. Neurobiol Dis 23: 260–72

Benarroch EE. 2018. Glutamatergic synaptic plasticity and dysfunction in Alzheimer disease: Emerging mechanisms. Neurology 91: 125–32

Boche D, Nicoll JAR. 2020. Invited Review - Understanding cause and effect in Alzheimer’s pathophysiology: Implications for clinical trials. Neuropathol Appl Neurobiol 46: 623–40

Cao W, Zheng H. 2018. Peripheral immune system in aging and Alzheimer’s disease. Mol Neurodegener 13: 51

Carlesimo GA, Piras F, Orfei MD, Iorio M, Caltagirone C, Spalletta G. 2015. Atrophy of presubiculum and subiculum is the earliest hippocampal anatomical marker of Alzheimer’s disease. Alzheimer’s & Dementia: Diagnosis, Assessment & Disease Monitoring 1: 24–32

Ceeraz S, Nowak EC, Noelle RJ. 2013. B7 family checkpoint regulators in immune regulation and disease. Trends Immunol 34: 556–63

Chishti MA, Yang DS, Janus C, Phinney AL, Horne P, et al. 2001. Early-onset amyloid deposition and cognitive deficits in transgenic mice expressing a double mutant form of amyloid precursor protein 695. J Biol Chem 276: 21562–70

Chittimalli K, Adkins S, Arora S, Singh J, Jarajapu YPR. 2024. An Investigation of the Inflammatory Landscape in the Brain and Bone Marrow of the APP/PS1 Mouse. J Alzheimers Dis Rep 8: 981–98

Cortes-Canteli M, Kruyer A, Fernandez-Nueda I, Marcos-Diaz A, Ceron C, et al. 2019. Long-Term Dabigatran Treatment Delays Alzheimer’s Disease Pathogenesis in the TgCRND8 Mouse Model. J Am Coll Cardiol 74: 1910–23

Coulson EJ. 2006. Does the p75 neurotrophin receptor mediate Abeta-induced toxicity in Alzheimer’s disease? J Neurochem 98: 654–60

Deinhardt K, Kim T, Spellman DS, Mains RE, Eipper BA, et al. 2011. Neuronal growth cone retraction relies on proneurotrophin receptor signaling through Rac. Sci Signal 4: ra82

Ding SL. 2013. Comparative anatomy of the prosubiculum, subiculum, presubiculum, postsubiculum, and parasubiculum in human, monkey, and rodent. J Comp Neurol 521: 4145–62

Dudal S, Krzywkowski P, Paquette J, Morissette C, Lacombe D, Tremblay P, Gervais F. 2004. Inflammation occurs early during the Abeta deposition process in TgCRND8 mice. Neurobiol Aging 25: 861–71

Ernst PB, Carvunis AR. 2018. Of mice, men and immunity: a case for evolutionary systems biology. Nat Immunol 19: 421–25

Farkas E, Luiten PG. 2001. Cerebral microvascular pathology in aging and Alzheimer’s disease. Prog Neurobiol 64: 575–611

Felger JC, Abe T, Kaunzner UW, Gottfried-Blackmore A, Gal-Toth J, et al. 2010. Brain dendritic cells in ischemic stroke: time course, activation state, and origin. Brain Behav Immun 24: 724–37

Genc K, Dona DL, Reder AT. 1997. Increased CD80(+) B cells in active multiple sclerosis and reversal by interferon beta-1b therapy. J Clin Invest 99: 2664–71

Giza JI, Kim J, Meyer HC, Anastasia A, Dincheva I, et al. 2018. The BDNF Val66Met Prodomain Disassembles Dendritic Spines Altering Fear Extinction Circuitry and Behavior. Neuron 99: 163–78.e6

Granger MW, Franko B, Taylor MW, Messier C, George-Hyslop PS, Bennett SA. 2016. A TgCRND8 Mouse Model of Alzheimer’s Disease Exhibits Sexual Dimorphisms in Behavioral Indices of Cognitive Reserve. J Alzheimers Dis 51: 757–73

Grubman A, Choo XY, Chew G, Ouyang JF, Sun G, et al. 2021. Transcriptional signature in microglia associated with Abeta plaque phagocytosis. Nat Commun 12: 3015

Heneka MT, Gauthier S, Chandekar SA, Hviid Hahn-Pedersen J, Bentsen MA, Zetterberg H. 2025. Neuroinflammatory fluid biomarkers in patients with Alzheimer’s disease: a systematic literature review. Mol Psychiatry 30: 2783–98

Irmady K, Jackman KA, Padow VA, Shahani N, Martin LA, et al. 2014. Mir-592 regulates the induction and cell death-promoting activity of p75NTR in neuronal ischemic injury. J Neurosci 34: 3419–28

Jolas T, Zhang XS, Zhang Q, Wong G, Del Vecchio R, Gold L, Priestley T. 2002. Long-term potentiation is increased in the CA1 area of the hippocampus of APP(swe/ind) CRND8 mice. Neurobiol Dis 11: 394–409

Kim J, Yoo ID, Lim J, Moon JS. 2024. Pathological phenotypes of astrocytes in Alzheimer’s disease. Exp Mol Med 56: 95–99

Kisler K, Nelson AR, Montagne A, Zlokovic BV. 2017. Cerebral blood flow regulation and neurovascular dysfunction in Alzheimer disease. Nat Rev Neurosci 18: 419–34

Knopman DS, Amieva H, Petersen RC, Chetelat G, Holtzman DM, et al. 2021. Alzheimer disease. Nat Rev Dis Primers 7: 33

Kokaia Z, Andsberg G, Martinez-Serrano A, Lindvall O. 1998. Focal cerebral ischemia in rats induces expression of p75 neurotrophin receptor in resistant striatal cholinergic neurons. Neuroscience 84: 1113–25

Lee HG, Lee JH, Flausino LE, Quintana FJ. 2023. Neuroinflammation: An astrocyte perspective. Sci Transl Med 15: eadi7828

Madsen LS, Parbo P, Ismail R, Gottrup H, Ostergaard L, Brooks DJ, Eskildsen SF. 2023. Capillary dysfunction correlates with cortical amyloid load in early Alzheimer’s disease. Neurobiol Aging 123: 1–9

Malik SC, Sozmen EG, Baeza-Raja B, Le Moan N, Akassoglou K, Schachtrup C. 2021. In vivo functions of p75(NTR): challenges and opportunities for an emerging therapeutic target. Trends Pharmacol Sci 42: 772–88

Marongiu R, Platholi J, Park L, Yu F, Sommer G, et al. 2025. Promotion of neuroinflammation in select hippocampal regions in a mouse model of perimenopausal Alzheimer’s disease. Front Mol Biosci 12: 1597130

Mathys H, Boix CA, Akay LA, Xia Z, Davila-Velderrain J, et al. 2024. Single-cell multiregion dissection of Alzheimer’s disease. Nature 632: 858–68

Meeker RB, Poulton W, Feng WH, Hudson L, Longo FM. 2012. Suppression of immunodeficiency virus-associated neural damage by the p75 neurotrophin receptor ligand, LM11A-31, in an in vitro feline model. J Neuroimmune Pharmacol 7: 388–400

Moonen S, Koper MJ, Van Schoor E, Schaeverbeke JM, Vandenberghe R, et al. 2023. Pyroptosis in Alzheimer’s disease: cell type-specific activation in microglia, astrocytes and neurons. Acta Neuropathol 145: 175–95

Morano NC, Smith RS, Danelon V, Schreiner R, Patel U, et al. 2022. Human immunomodulatory ligand B7-1 mediates synaptic remodeling via the p75 neurotrophin receptor. J Clin Invest 132

Morris GP, Foster CG, Courtney JM, Collins JM, Cashion JM, et al. 2023. Microglia directly associate with pericytes in the central nervous system. Glia 71: 1847–69

Nortley R, Korte N, Izquierdo P, Hirunpattarasilp C, Mishra A, et al. 2019. Amyloid beta oligomers constrict human capillaries in Alzheimer’s disease via signaling to pericytes. Science 365

Patnaik A, Zagrebelsky M, Korte M, Holz A. 2020. Signaling via the p75 neurotrophin receptor facilitates amyloid-beta-induced dendritic spine pathology. Sci Rep 10: 13322

Perini G, Della-Bianca V, Politi V, Della Valle G, Dal-Pra I, Rossi F, Armato U. 2002. Role of p75 neurotrophin receptor in the neurotoxicity by beta-amyloid peptides and synergistic effect of inflammatory cytokines. J Exp Med 195: 907–18

Platholi J, Marongiu R, Park L, Yu F, Sommer G, et al. 2023. Hippocampal glial inflammatory markers are differentially altered in a novel mouse model of perimenopausal cerebral amyloid angiopathy. Front Aging Neurosci 15: 1280218

Religa P, Cao R, Religa D, Xue Y, Bogdanovic N, et al. 2013. VEGF significantly restores impaired memory behavior in Alzheimer’s mice by improvement of vascular survival. Sci Rep 3: 2053

Rogers J, Luber-Narod J, Styren SD, Civin WH. 1988. Expression of immune system-associated antigens by cells of the human central nervous system: relationship to the pathology of Alzheimer’s disease. Neurobiol Aging 9: 339–49

Roy DS, Kitamura T, Okuyama T, Ogawa SK, Sun C, et al. 2017. Distinct Neural Circuits for the Formation and Retrieval of Episodic Memories. Cell 170: 1000–12 e19

Santisteban MM, Iadecola C. 2025. The pathobiology of neurovascular aging. Neuron 113: 49–70

Schildberg FA, Klein SR, Freeman GJ, Sharpe AH. 2016. Coinhibitory Pathways in the B7-CD28 Ligand-Receptor Family. Immunity 44: 955–72

Sebastiani A, Golz C, Werner C, Schafer MK, Engelhard K, Thal SC. 2015. Proneurotrophin Binding to P75 Neurotrophin Receptor (P75ntr) Is Essential for Brain Lesion Formation and Functional Impairment after Experimental Traumatic Brain Injury. J Neurotrauma 32: 1599–607

Siao CJ, Lorentz CU, Kermani P, Marinic T, Carter J, et al. 2012. ProNGF, a cytokine induced after myocardial infarction in humans, targets pericytes to promote microvascular damage and activation. J Exp Med 209: 2291–305

Spires-Jones TL, Hyman BT. 2014. The intersection of amyloid beta and tau at synapses in Alzheimer’s disease. Neuron 82: 756–71

Steele JW, Brautigam H, Short JA, Sowa A, Shi M, et al. 2014. Early fear memory defects are associated with altered synaptic plasticity and molecular architecture in the TgCRND8 Alzheimer’s disease mouse model. J Comp Neurol 522: 2319–35

Sugiura D, Maruhashi T, Okazaki IM, Shimizu K, Maeda TK, Takemoto T, Okazaki T. 2019. Restriction of PD-1 function by cis-PD-L1/CD80 interactions is required for optimal T cell responses. Science 364: 558–66

Thome AD, Faridar A, Beers DR, Thonhoff JR, Zhao W, et al. 2018. Functional alterations of myeloid cells during the course of Alzheimer’s disease. Mol Neurodegener 13: 61

Turtzo LC, Lescher J, Janes L, Dean DD, Budde MD, Frank JA. 2014. Macrophagic and microglial responses after focal traumatic brain injury in the female rat. J Neuroinflammation 11: 82

Uemura MT, Maki T, Ihara M, Lee VMY, Trojanowski JQ. 2020. Brain Microvascular Pericytes in Vascular Cognitive Impairment and Dementia. Front Aging Neurosci 12: 80

Vara-Perez M, Movahedi K. 2025. Border-associated macrophages as gatekeepers of brain homeostasis and immunity. Immunity 58: 1085–100

West SM, Deng XA. 2019. Considering B7-CD28 as a family through sequence and structure. Exp Biol Med (Maywood) 244: 1577–83

Windhagen A, Newcombe J, Dangond F, Strand C, Woodroofe MN, Cuzner ML, Hafler DA. 1995. Expression of costimulatory molecules B7-1 (CD80), B7-2 (CD86), and interleukin 12 cytokine in multiple sclerosis lesions. J Exp Med 182: 1985–96

Woo NH, Teng HK, Siao CJ, Chiaruttini C, Pang PT, et al. 2005. Activation of p75NTR by proBDNF facilitates hippocampal long-term depression. Nat Neurosci 8: 1069–77

Wykes R, Kalmbach A, Eliava M, Waters J. 2012. Changes in the physiology of CA1 hippocampal pyramidal neurons in preplaque CRND8 mice. Neurobiol Aging 33: 1609–23

Yamashita T, Tucker KL, Barde YA. 1999. Neurotrophin binding to the p75 receptor modulates Rho activity and axonal outgrowth. Neuron 24: 585–93

Yan P, Kim KW, Xiao Q, Ma X, Czerniewski LR, et al. 2022. Peripheral monocyte-derived cells counter amyloid plaque pathogenesis in a mouse model of Alzheimer’s disease. J Clin Invest 132

Yang AC, Vest RT, Kern F, Lee DP, Agam M, et al. 2022. A human brain vascular atlas reveals diverse mediators of Alzheimer’s risk. Nature 603: 885–92

Yang J, Harte-Hargrove LC, Siao CJ, Marinic T, Clarke R, et al. 2014. proBDNF negatively regulates neuronal remodeling, synaptic transmission, and synaptic plasticity in hippocampus. Cell Rep 7: 796–806

Zenaro E, Piacentino G, Constantin G. 2017. The blood-brain barrier in Alzheimer’s disease. Neurobiol Dis 107: 41–56

Zhou Y, Song WM, Andhey PS, Swain A, Levy T, et al. 2020. Human and mouse single-nucleus transcriptomics reveal TREM2-dependent and TREM2-independent cellular responses in Alzheimer’s disease. Nat Med 26: 131–42

Ziar R, Tesar PJ, Clayton BLL. 2025. Astrocyte and oligodendrocyte pathology in Alzheimer’s disease. Neurotherapeutics 22: e00540

Zlokovic BV. 2011. Neurovascular pathways to neurodegeneration in Alzheimer’s disease and other disorders. Nat Rev Neurosci 12: 723–38

Zotova E, Bharambe V, Cheaveau M, Morgan W, Holmes C, et al. 2013. Inflammatory components in human Alzheimer’s disease and after active amyloid-beta42 immunization. Brain 136: 2677–96

