## Supplementary material for "Expression of the human immunomodulatory protein, human B7-1 (CD80), accelerates neuroinflammation, synaptic loss, microvascular instability and lethality in a murine model of Alzheimer’s Disease": Supp. Figures+Legends

**A**

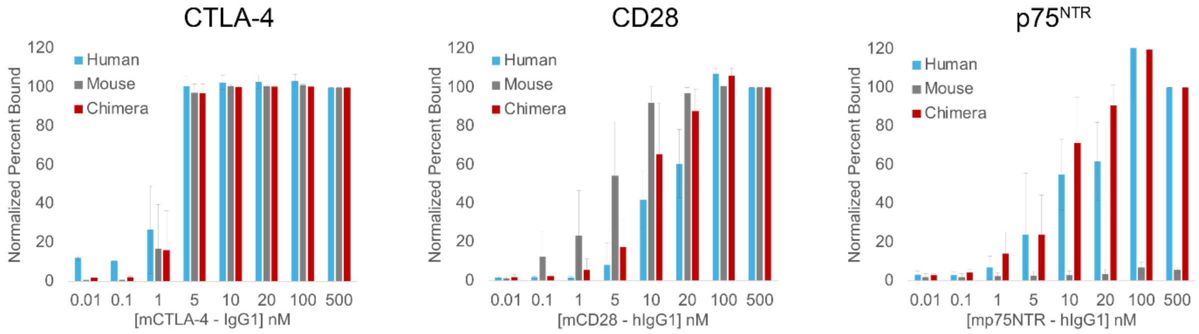


**B**

| | EC50 CTLA-4 (nM) | EC50 CD28 (nM) | EC50 p75 NTR (nM) |
|---|---|---|---|
| Human | 1.6 ± 0.7 | 15.0 ± 3.9 | 13.6 ± 8.7 |
| Mouse | 1.7 ± 0.6 | 5.0 ± 2.2 | X |
| Chimera | 1.8 ± 0.6 | 8.6 ± 1.8 | 8.6 ± 2.3 |

Supp Figure 1

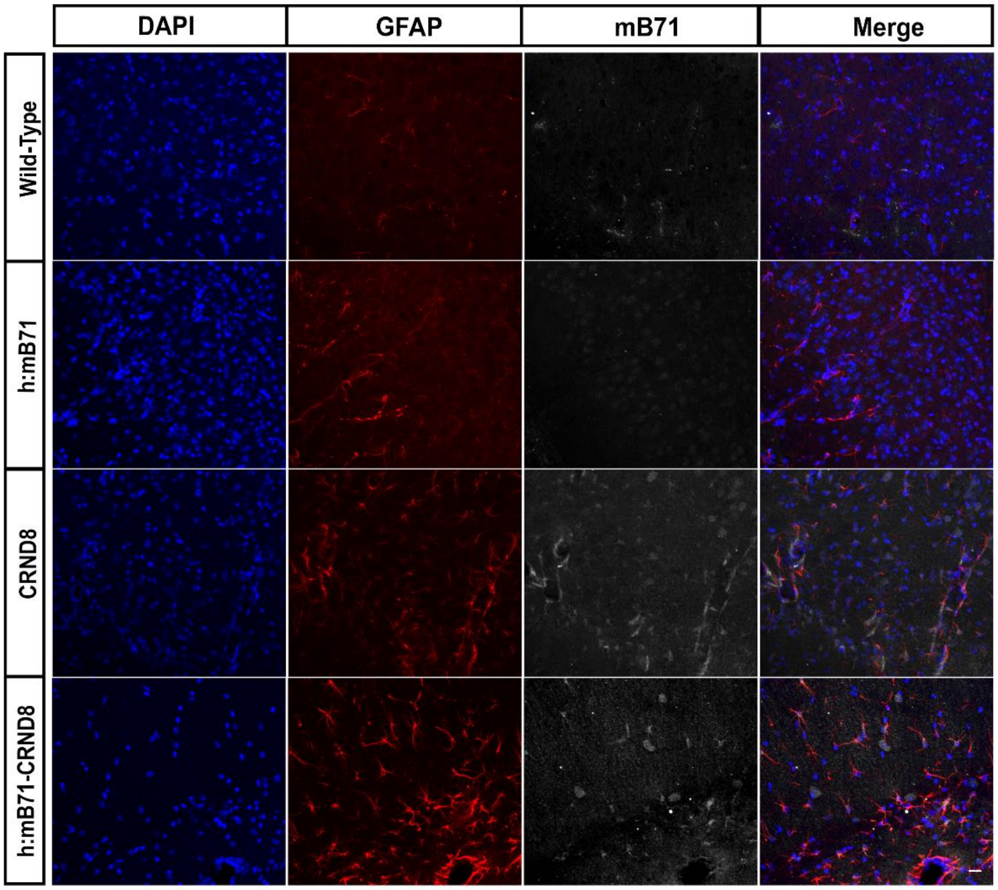


Supp Figure 2

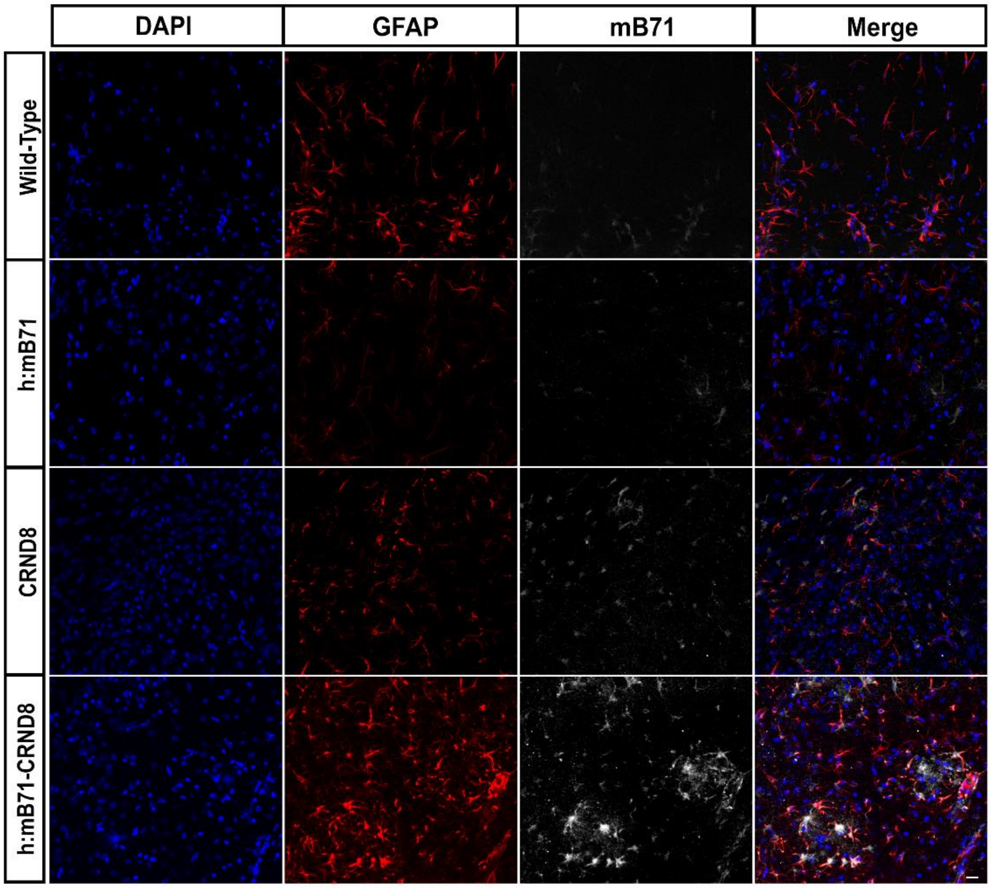


Supp Figure 3

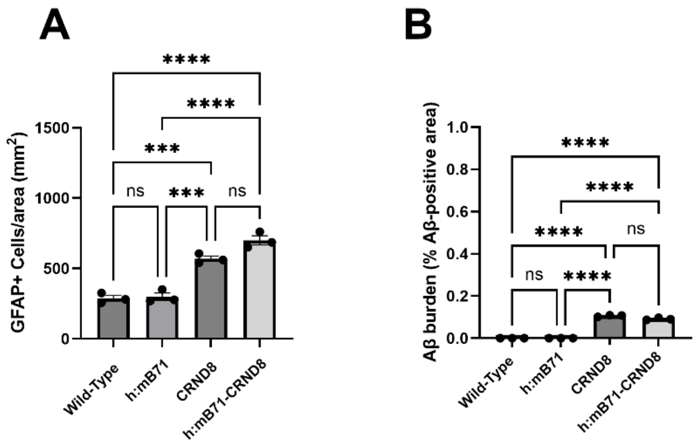


Supp Figure 4

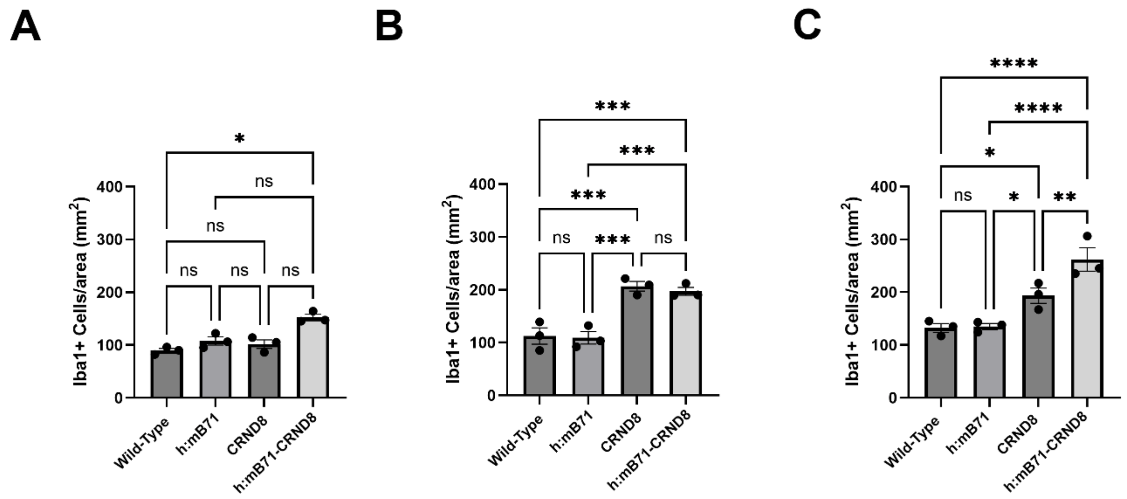


Supp Figure 5

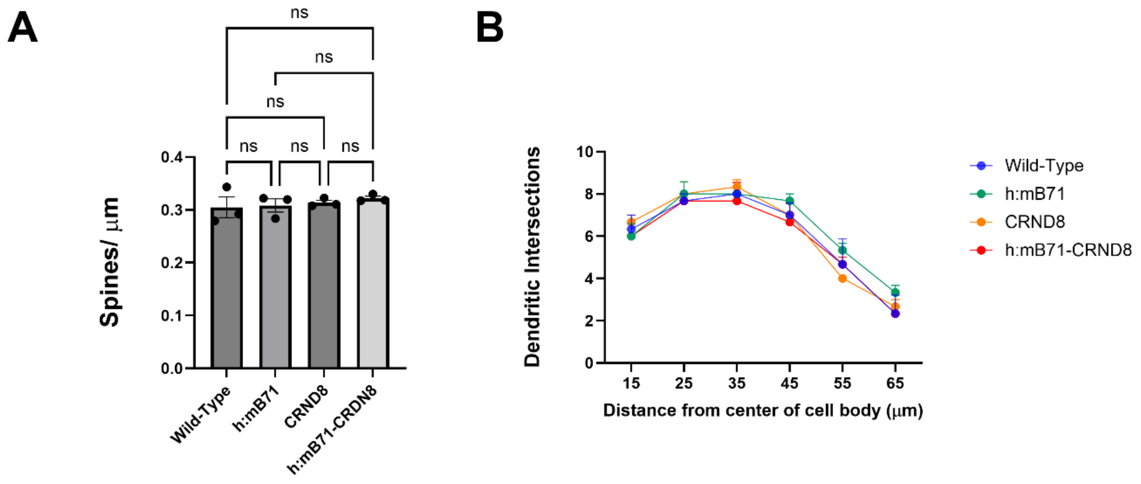


Supp Figure 6

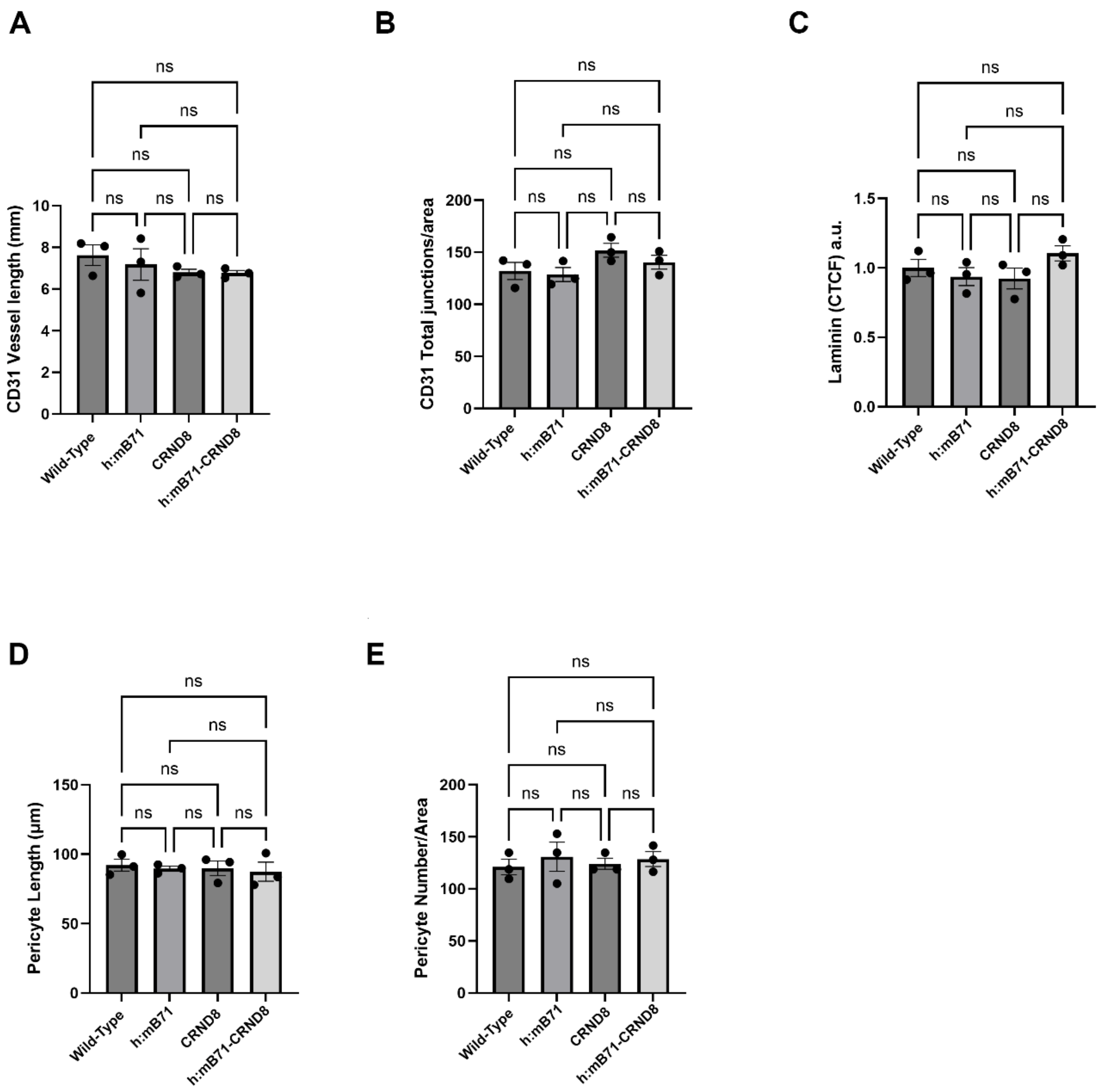


Supp Figure 7

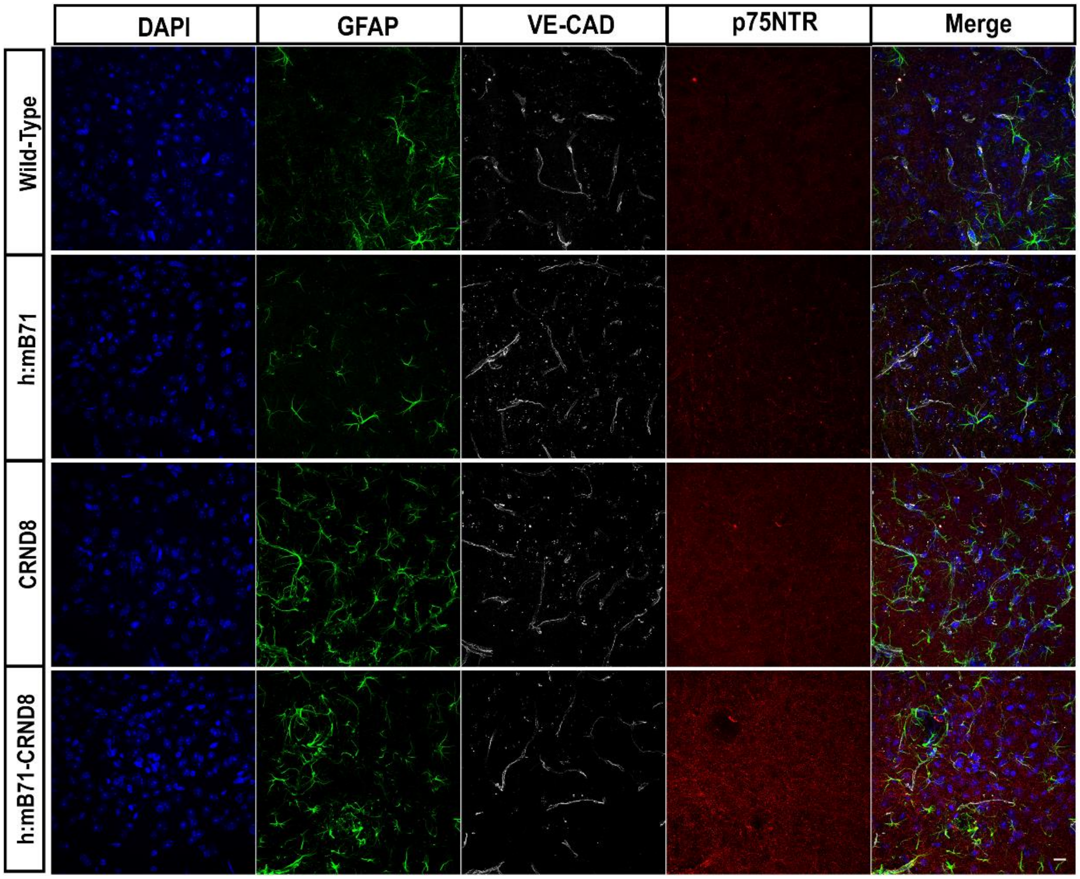


Supp Figure 8

### Supplementary Figure Legends

**Supplementary Figure 1**. *Extended data and calculated EC50 values for the titrations of mCTLA-4, mCD28, and mp75NTR hIgG1 proteins against cells expressing human, mouse or chimera B7-1.* A. Data shows bar graph representations of the titration experiment described in Figure 1A. B. EC50 values were calculated by fitting the titration data shown in Figure 1A to a standard single-site binding equation with variable slope using non-linear regression in Graphpad Prism ($Y=B_{min} + (B_{max}-B_{min})/(1+10^{((LogEC50-X)*HillSlope)})$). These values highlight the similarity in binding between the B7-1 constructs tested, especially between human and the h:m chimera.

**Supplementary Figure 2.** *mB71 expression in mice of 3 months of age***.** Representative confocal images of WT, h:mB71, CRND8 and h:mB71-CRND8 mice showing DAPI stained nuclei, GFAP and mB71 expression. mB71 protein is upregulated in wt/CRND8 and h:mB71-CRND8 in GFAP+ cells, and more rarely in GFAP- cells. Scale bar 10 µm.

**Supplementary Figure 3.** *mB71 expression in mice of 4 months of age***.** Representative confocal images of WT, h:mB71, CRND8 and h:mB71-CRND8 mice showing DAPI stained nuclei, GFAP and mB71 expression. mB71 protein is upregulated in CRND8 and h:mB71-CRND8 in GFAP+ cells and more rarely in GFA- cells . Scale bar 10 µm

**Supplementary Figure 4.** *Induction of GFAP+cells, and Aβ in 2 month old wt/CRND8 and h:mB7-1/CRND8 mice* Quantification of confocal images of wt/wt, h:mB7-1/wt, wt/CRND8, h:mB7-1/CRND8 mice immunostained for GFAP and beta amyoid. With Aβ overexpression, GFAP was comparably induced in wt/CRND8, h:mB7-1/CRND8 mice.

**Supplementary Figure 5.** *Iba1+ cells in mice of 2, 3 and 4months of age in the dSubiculum increases*. Quantitative analysis of confocal images immunostained for Iba1. (A) Number of Iba1+ cells/area in wt/wt, h:mB71/wt, wt/CRND8 and h:mB71/CRND8 mice at 2mo. (B) Number of Iba1+ cells/area in in wt/wt, h:mB71/wt, wt/CRND8 and h:mB71/CRND8 mice at 3mo. (C) Number of Iba1+ cells/area in in wt/wt, h:mB71/wt, wt/CRND8 and h:mB71/CRND8 mice at 4mo. In (A) *p=0.0231., In (B) ***p=0.0009 (wt/wt vs h:mB71/CRND8), ***p=0.0005 (h:mB71/wt vs h:mB71/CRND8), ***p=0.0002 (wt/wt vs wt/CRND8), ***p=0.0001 (h:mB71/wt vs wt/CRND8). In (C) ****<0.0001 (wt/wt vs h:mB71/CRND8), ***p=0.0001 (h:mB71/wt vs h:mB71/CRND8), *p=0.0293 (wt/wt vs wt/CRND8), ***p=0.0425 (h:mB71/wt vs wt/CRND8), **p=0.0097 (wt/CRND8 vs h:mB71/CRND8).

**Supplementary Figure 6.** *Golgi analysis of mice at 2months of age. No significant differences were found in dendritic spine density and basal dendritic arborization among the genotypes.* (A) Quantification of Golgi stained images revealed no statistic differences among the genotypes in dendritic spines density or (B) using Sholl analysis, in basal dendritic intersections. Data shown are mean +/- SEM. In (A) $p>0.05$, one-way ANOVA followed by post hoc Tukey test). N=3.

**Supplementary Figure 7.** *Analysis of the Vasculature of mice at 2 months of age***.** *No significant differences were found vessel length, vessel junction in the microvasculature,or in laminin expression and pericyte number and length.* Quantitative analysis of confocal images from in wt/wt, h:mB71/wt, wt/CRND8 and h:mB71/CRND8 mice (A) Vessel length, (B) Total number of junctions, (C) Laminin protein levels, (D) Pericyte length and (E) Pericyte number per area. n.s. not significant. One-way ANOVA followed by post hoc Tukey test. N=3.
